# Electroconvulsive therapy generates a postictal wave of spreading depolarization in mice and humans

**DOI:** 10.1101/2024.10.31.621357

**Authors:** Zachary P Rosenthal, Joseph B. Majeski, Ala Somarowthu, Davin K Quinn, Britta E. Lindquist, Mary E. Putt, Antoneta Karaj, Chris G Favilla, Wesley B. Baker, Golkoo Hosseini, Jenny P Rodriguez, Mario A Cristancho, Yvette I Sheline, C. William Shuttleworth, Christopher C. Abbott, Arjun G Yodh, Ethan M Goldberg

## Abstract

Electroconvulsive therapy (ECT) is a fast-acting, highly effective, and safe treatment for medication-resistant depression. Historically, the clinical benefits of ECT have been attributed to generating a controlled seizure; however, the underlying neurobiology is understudied and unresolved. Using optical neuroimaging of neural activity and hemodynamics in a mouse model of ECT, we demonstrated that a second brain event follows seizure: cortical spreading depolarization (CSD). We found that ECT pulse parameters and electrode configuration directly shaped the wave dynamics of seizure and subsequent CSD. To translate these findings to human patients, we used non-invasive diffuse optical monitoring of cerebral blood flow and oxygenation during routine ECT treatments. We observed that human brains reliably generate hyperemic waves after ECT seizure which are highly consistent with CSD. These results challenge a long-held assumption that seizure is the primary outcome of ECT and point to new opportunities for optimizing ECT stimulation parameters and treatment outcomes.

## Introduction

Electroconvulsive therapy (ECT) is a life-saving intervention for medication-resistant depression. Treatment consists of electrical pulses delivered to the brain to elicit a controlled (∼30-90 second) electrographic seizure while under general anesthesia and muscle relaxants to minimize physical movement. Nearly a century since it was discovered, ECT remains the most clinically effective treatment for severe depression^1^. A typical index course of six to twelve ECT treatments achieves rapid symptom improvement in 60-80% of patients, reducing suicide risk by 50% compared to matched controls who do not receive ECT^2,3^. ECT is also highly effective in mania, psychosis, Parkinson’s disease, catatonia, and even status epilepticus – particularly for those with the most severe, medication-resistant symptoms. The procedure is safe and generally well-tolerated, including in pediatric, geriatric, and pregnant populations. Potential risks, such as cognitive slowing or memory impairment, are typically modest compared to improvement in neuropsychiatric symptoms, and resolve when treatment is discontinued. Unfortunately, the proven efficacy and safety of ECT have been overshadowed by stigma and negative portrayal in popular media. As a result, contemporary research on ECT is limited, and the treatment is underutilized^4^. Much could be learned from ECT and the basic physiology underlying its rapid-acting efficacy across a varied range of brain disorders, particularly when modern pharmacology has failed.

For the last eight decades, it has been assumed that seizure is necessary for ECT to elicit clinical improvement^3^. However, not all ECT-induced seizures are therapeutic, and conversely, electrical stimulation below the seizure threshold may also be an effective treatment^5,6^. The role of seizure in the therapeutic mechanism of ECT is thus uncertain. An important clue to this mechanism may be that ECT induces lasting inhibitory plasticity in brain activity, dampening cortical response to stimulation and reliably reducing propensity for future seizures (i.e., raising the seizure threshold)^7–11^. It has been hypothesized that ECT may elicit inhibitory plasticity through the process of seizure suppression^12^, but few studies have explored the cellular, circuit, or network level mechanisms of brain inhibition after ECT.

During ECT, brain activity is typically monitored with a minimal scalp electroencephalography (EEG) montage consisting of two leads on the forehead to measure seizure quality, duration and post-ictal suppression in each hemisphere. However, quantitative seizure metrics from EEG monitoring have shown limited predictive value for ECT clinical outcomes^5–7,13–16^. A great need exists for more robust brain activity biomarkers that can predict treatment efficacy, detect side effects, or guide selection of stimulation parameters. Because ECT electrical pulses saturate the scalp EEG signal, no study has ever measured brain activity during pulse delivery in human patients or animal models. Computational models have helped predict the brain electric field evoked by a single electrical pulse^5,17–19^, but there are likely opportunities to further optimize stimulation parameters and improve clinical outcomes using empirical measurements of brain activity during treatment. For example, a seminal randomized controlled trial showed that the choice of two electrode spatial configurations (right unilateral vs. bitemporal), as well as two pulse durations (0.3 vs. 0.5 ms), significantly modulates the clinical efficacy and side effects of ECT^20,21^. The remaining stimulation parameter space is vast, mostly untested, and with unknown effects on brain activity and clinical outcomes^20^. This includes infinite combinations of pulse properties (duration, frequency, total count, current, polarity, waveform) and electrode spatial configurations. Only three electrode placements have been tested clinically – right unilateral, bitemporal, and bifrontal. There are likely alternative stimulation parameters that would provide superior symptom reduction and lower risk of side effects compared to the current standard-of-care configurations and titration algorithms.

To bridge these knowledge gaps, we employed optical neuroimaging to record brain activity during ECT in both rodent models and human patients. Our data demonstrate that a second brain event follows ECT-induced seizure: cortical spreading depolarization (CSD). In mice, we show that clinically relevant choices of electrode placements and pulse parameters directly modulate the spatial and temporal properties of both seizure and subsequent CSD. We then translate these findings from mice to humans. Using non-invasive diffuse optical monitoring of brain tissue oxygen saturation and blood flow in patients receiving ECT, we find evidence for post-ictal hyperemic waves that are highly consistent with CSD. These discoveries have important implications for ECT’s mechanism of action and for optimizing stimulation to target specific brain outcomes.

## Methods

### Mouse model

All procedures described below were approved by the Institutional Animal Care and Use Committee at the Children’s Hospital of Philadelphia in compliance with AAALAC guidelines. Mice were raised in standard cages in a double barrier mouse facility with a 12-hr/12-hr light/dark cycle and *ad libitum* access to food and water. All experiments used n=10 eight-week-old mice (male and female) hemizygous for *Thy1-jRGECO1a* (JAX 030525) on a C57BL/6J background, to enable optical imaging of the fluorescent calcium sensor protein jRGECO1a. Pups were genotyped by PCR prior to experiments to confirm the presence of the *Thy1-jRGECO1a* transgene, using the forward primer 5’- ACAGAATCCAAGTCGGAACTC-3’ and reverse primer 5’- CCTATAGCTCTGACTGCGTGAC-3’.

### Cranial window and electrode implantation

Mice were treated preoperatively with subcutaneous buprenorphine-SR (1.0 mg/kg), meloxicam (5.0 mg/kg) and cefazolin (500 mg/kg). Mice were anesthetized with isoflurane (3% induction, 1.5% maintenance). Body temperature was maintained via heating pad. The scalp was shaved, sterilized with alcohol and betadine, incised at midline, and retracted to expose the dorsal skull. Five brass electrode pins with 1.78mm diameter flanges (DigiKey 4443-0-00-15-00-00-03-0) were attached to the intact skull surface using a thin layer of silver conductive epoxy (MG Chemicals 8331D). Electrodes were stereotactically centered relative to bregma (see **Fig. 1a**; #1, frontal: X = 0.00 mm, Y = 4.09 mm; #2 and #3, temporo-parietal: X = ±3.40 mm, Y = 0.47 mm; #4 and #5, occipital: X = ±4.69 mm, Y = -4.89 mm). A custom steel headbar was attached to the posterior skull and a cranial window was formed with optically clear dental cement (C&B-Metabond, Parkell Inc., Edgewood, NY). The window was rendered transparent and hardened with clear UV-cure gel nail polish. Animals were allowed to recover from surgery for at least one week prior to imaging.

**Fig. 1.**
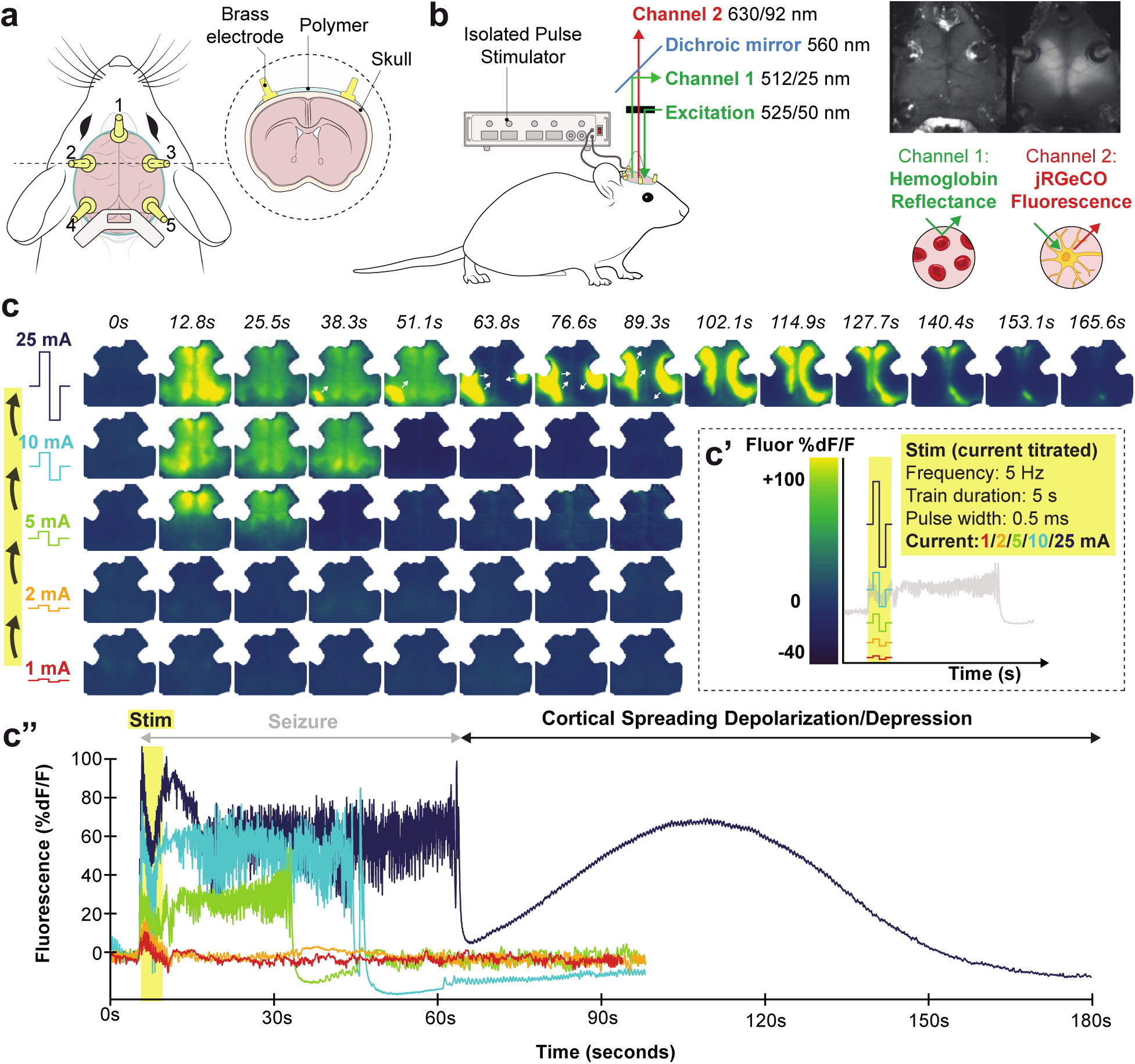
Optical neuroimaging in a mouse model of ECT demonstrates a current threshold where seizure is followed by cortical spreading depolarization. **a**, Schematic of mouse cranial window with optical access to the dorsal cortex. Within each window, five brass electrodes were attached to the intact skull overlying frontal (#1), temporo-parietal (#2, 3), and occipital (#4, 5) cortices. **b**, Green-filtered illumination (525/50 nm) was projected through the microscope objective onto the brain; reflected and emitted light from the brain was separated by a 560 nm dichroic mirror into two spatially co-registered channels that were bandpass filtered for isosbestic point hemoglobin reflectance of green illumination (512/25 nm) or jRGECO1a red fluorescence emission (630/92 nm). Electrical stimulation was delivered via pairs of brass electrode pins using an isolated pulse stimulator. See **Extended Data Fig. 1** and **Supplemental Methods**. **c**, Example video frames (from **Extended Data Video 1**) of neuronal activity (jRGECO1a relative fluorescence %dF/F) in an individual mouse being stimulated with 5 Hz pulses to electrodes 1 and 4. Stimulation was titrated in successive rounds with increasing current (**c**’, note rainbow color map corresponds to current titration); in each round we monitored brain activity for 5 seconds of baseline, during stimulation, and then for 90 seconds (or the resumption of baseline slow wave dynamics.). Data from each round of stimulation in the current titration are depicted in still-frame maps of cortical activity in (**c**) and time series averaged within a right hemisphere ROI (**c**”).

### Widefield imaging system

Widefield imaging of neuronal dynamics (jRGECO1a fluorescence) and cerebral hemodynamics (optical intrinsic signal, OIS) was performed using a modified version of a previous method^22^, on a Leica M205FA stereoscope fitted with a 0.63× objective to capture the dorsal convexity of the cortex (a ∼1 cm^2^ area, binned to 128×128 pixels, 0.078 mm pixel-width). System optical spectra are depicted in **Extended Data Fig. 1**. White light LED illumination (X-Cite XYLIS) was bandpass filtered to green (525/50 nm, Chroma) and directed through the objective at the cortex. Light reflected and emitted from the brain was then separated into two channels using a 560 nm dichroic and image-splitting optics (Hamamatsu W-VIEW GEMINI). The OIS channel was filtered with a 512/25 nm bandpass and attenuated with a 5% transmission neutral density filter (Chroma), optimized to capture light reflectance at the isosbestic point of hemoglobin to approximate total hemoglobin concentration. The jRGECO1a red fluorescence channel was filtered with a 630/92 nm bandpass filter. Both channels were detected side-by-side and spatially co-registered on a single CMOS camera (Hamamatsu ORCA-Flash4.0 V3). An exposure time of 50 ms was used, achieving a sampling rate of 15.7 Hz.

### Combined imaging and electroconvulsive therapy in mice

Mice were monitored with head-fixed widefield imaging while under dexmedetomidine anesthesia (0.5 mg/kg, IP), which reliably achieved a plane of sedation in ∼10 minutes with loss of response to toe-pinch and emergence of ∼3 Hz anteroposterior global slow waves that are characteristic of anesthesia^23,24^. ECT pulses were delivered to pairs of cranial electrodes using constant current pulse trains (described below) with an isolated pulse stimulator (A-M Systems 4100). At the end of each recording session, anesthesia was reversed with atipamezole (0.5 mg/kg). Similar to prior reports, seizures were noted to elicit ∼1-60 seconds of high-amplitude, aperiodic neuronal fluorescence activity globally across the cortex, as well as tonic-clonic limb and tail movements. Several pilot titrations were also conducted using etomidate as well as ketamine/xylazine anesthesia, which likewise demonstrated CSD after ECT seizure (data not shown); dexmedetomidine was ultimately selected because of its hemodynamic safety^25^, reversibility, and favorable pharmacodynamic profile as an alpha agonist, avoiding acting directly on GABAergic or glutamatergic currents.

To broadly survey the stimulation parameter space, n=38 ECT stimulus titrations were performed, each testing a different combination of electrode spatial configuration (5 electrodes, 10 possible pairs), frequency (5, 10, 25, 50, 100 Hz) and pulse counts (25, 50, 100 pulses); as shown in **Extended Data Table 1**. All stimulation trials used constant-current, bipolar square waves of 0.5 ms pulse width, similar to routine brief pulse ECT in humans. Within each titration, stimulation was delivered first at 1 mA current and then successively up-titrated to 2 mA, 5 mA, 10 mA, and 25 mA, akin to the individualized low amplitude seizure therapy (ILAST) strategy of ECT^26^. At each current step, brain activity was monitored at baseline for 5 seconds, during stimulation, and then for at least 90s, or until the sustained return of baseline anteroposterior slow waves from anesthesia, before being restimulated at the next highest current step. At the current threshold when seizure elicited CSD, recordings were continued for 10 minutes, and the titration was terminated. Titration sessions within individual mice were spaced at least 1 week apart to facilitate washout of acute effects of stimulation.

To control for effects of resampling the same individuals across titrations with different conditions, mixed effects modeling adjusted for mouse ID. Stimulation conditions were partially randomized and balanced across n=10 mice such that each mouse was treated with multiple frequencies and electrode configurations in varying session order. For any given electrode spatial configuration, multiple frequencies were tested; for each frequency, a mixture of unilateral and bilateral electrode configurations were tested. In n=4 titrations, animals were restimulated one more time to elicit a second CSD – these four secondary CSDs exhibited similar intrinsic amplitude and duration to initial CSDs, and were thus pooled into **Figs. 2a, 2g**, and **2h** (n=42). Secondary CSDs were excluded from analysis in **Figs. 2f** and **3**. One titration was excluded from analysis because of inadequate head-fixation leading to motion artifact that precluded accurate quantification; it was however visually scored as a right-sided CSD and pooled into **Fig. 2f** for completeness (n=39).

**Fig. 2.**
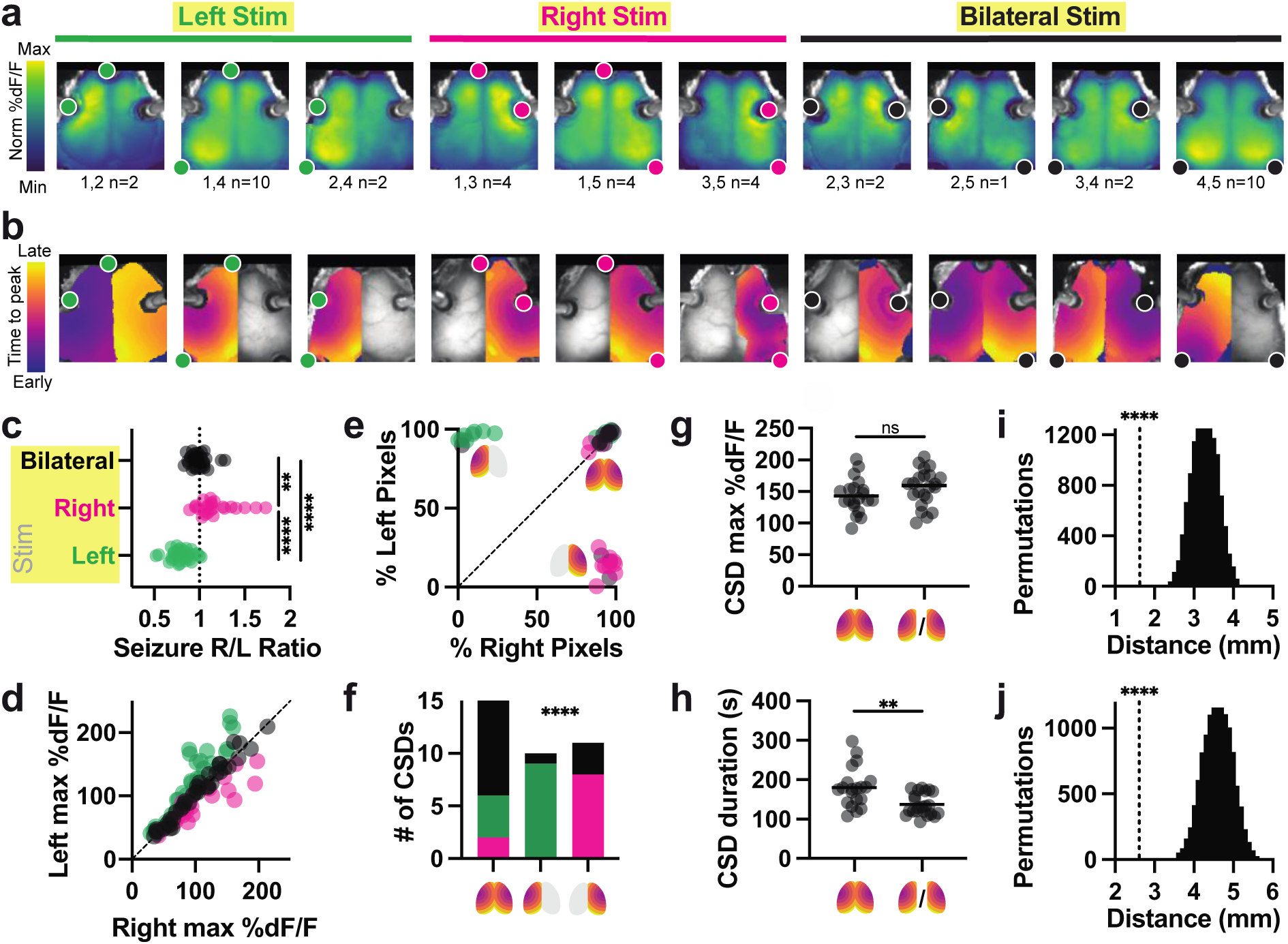
Electrode configuration shapes the spatial topography of seizure and CSD. **a,** Group average maps of seizure amplitude at each pixel (%dF/F). Each panel represents one of 10 possible pairs of 5 electrodes, averaged across multiple stimulation trials (electrode pair and number of trials in each average below, from n=42 end-titration seizures in n=10 mice). Fluorescence colormap is normalized to each plot’s min and max; absolute values of fluorescence are shown in **c** and **d**. Stimulated electrode pairs are depicted as colored dots. Note seizures tended to localize near stimulation electrodes. **b,** Example individual map of post-ictal CSD trajectory for each electrode configuration, depicting time-to-CSD-peak at each pixel. Note that some CSDs were recruited bilaterally to the entirety of both hemispheres, while others were recruited unilaterally to one hemisphere. Bilateral CSDs typically had some degree of temporal lag between the two hemispheres. Colormaps are normalized to the earliest and latest pixel, with absolute values of overall event durations shown in h. **c,** Asymmetry index for all detected seizures across all titrations (see Methods) evoked by right (R, n=25), left (L, n=34) and bilateral (BL, n=38) electrode configurations. Asymmetry index was calculated as the ratio of the left:right hemisphere maximum seizure amplitude averaged across pixels within each hemisphere. Kruskal-Wallis test with Dunn’s multiple comparison correction L vs. R (p**** < 0.0001), L vs BL (p****< 0.0001), R vs BL (p**=0.0026); Kruskal-Wallis statistic 61.83. **d,** Seizure amplitude (fluorescence, (%dF/F) values averaged within the right (x-axis) and left (y-axis) hemispheres for each event. A range of frequency and current conditions were tested (see Fig. 3), producing a wide range of fluorescence intensities. Electrode configuration for each seizure is color coded (L green, R magenta, BL Black). **e,** Percentage of pixels within the left hemisphere and right hemisphere where CSD was detected, from n=42 recorded events. Icons depict right-sided, left-sided, and bilateral CSD event types. **f,** Distribution of n=39 CSD outcomes resulting from right (magenta), left (green) or bilateral (black) stimulation. Fisher’s exact test with null hypothesis that the distributions of electrode configurations are equal in all three CSD outcomes (p < 0.0001 for 3x3 interaction). **g,** Average amplitude (%dF/F) for bilateral and unilateral CSDs (selecting whichever hemisphere had the largest activation). CSD amplitude was similar whether it occurred in one hemisphere or both, via two-tailed Mann-Whitney test, p=0.3306 (U=171, n=42). **h,** Overall duration of bilateral CSDs was slightly longer than that of unilateral CSDs, two-tailed Mann-Whitney test p**=0.0011 (U= 94, n=42). This was due to asymmetry in the start times for each hemisphere, illustrated in bilateral CSDs shown in b. **i,** Average distance (millimeters) of seizure peak pixel to nearest stimulation electrode (n=42 end-titration seizures), compared to 10,000 permutations of shuffled data, using a two-tailed t-test with Bonferroni corrected p****< 0.0001. **j,** Average distance (millimeters) of seizure peak pixel to CSD initiation pixel. Same statistical approach as i.

**Fig. 3.**
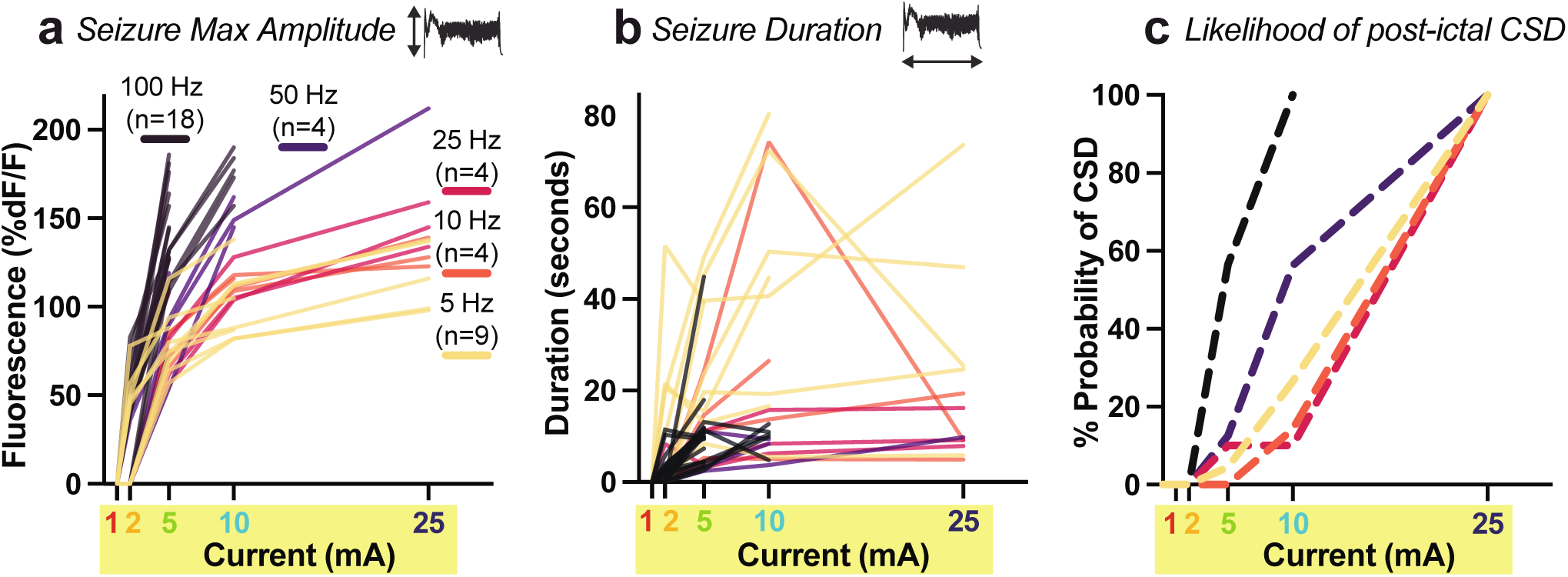
Pulse current and frequency modulate evoked seizure amplitude and duration, as well as subsequent cortical spreading depolarization. Each line in **a** and **b** represents an individual mouse being serially stimulated at a fixed pulse frequency (5, 10, 25, 50, or 100 Hz, yellow-orange-red-black color coding). Mice were first stimulated at 1 mA current, seizure was monitored until return to baseline, and then mice were restimulated in increasing in steps to 2, 5, 10, and 25 mA (rainbow color coding) until the evoked seizure was sufficient to trigger CSD. Individual lines for each mouse titration in **a** and **b** thus end at the current step that elicited CSD. The x-axis in **a**-**c** corresponds to the current step in the titration, while the y-axes measure three different brain event outcomes. **a,** Seizure amplitude (y-axis, fluorescence intensity %dF/F) averaged across the whole cortex as a function of current step (x-axis, mA) for each mouse titration (individual lines) at one of five frequencies (yellow-orange-red-black color coding). Data analyzed using mixed effects modeling to account for repeated trials on individual animals (see Methods). n is the number of individual mouse titration trials tested at each frequency. **b,** Same seizure titrations as in a) but with y-axis measuring seizure duration (seconds). Note inversion of the order of frequency groups relative to a. **c,** Cumulative probability of experiencing CSD after seizure at each current step within each frequency condition. Each line represents a group summary for the fixed frequency conditions 5, 10, 25, 50 and 100 Hz. See main text for results of statistical analysis.

### Mouse widefield optical imaging signal processing

Data were processed using an open-source widefield optical imaging toolkit^27^. Imaging data were converted into two-channel tiff files. A binary brain mask was manually drawn in MATLAB for each recording; all subsequent analyses were performed on pixels labeled as brain. Image sequences from each mouse (as well as the brain mask for each mouse) were affine-transformed to Paxinos atlas space using the positions of bregma and lambda^28^. A standardized mask of the stimulation electrodes was approximated using Paxinos coordinates (see seizure to electrode distance permutation analysis below). Right and left hemispheric divisions were segmented using a straight line drawn through midline of each image. A dark image with no illumination was subtracted from all frames, and then data were spatially smoothed with a Gaussian filter. The jRGECO1a calcium-sensor fluorescence signal (%dF/F) and total hemoglobin OIS signal (%dA/A) were calculated at each pixel by subtracting and dividing the 20^th^ percentile value from the first 4 seconds of baseline (pre-stimulation) raw signal from each recording. The total hemoglobin OIS signal was multiplied by -1 (absorbance) so that positive sign changes indicate increased blood volume/total hemoglobin concentration. Red-shifted fluorophores have significantly reduced hemoglobin absorption artifact compared to green fluorophores^22^. Given that fluorescence %dF/F event peaks were more than an order of magnitude larger than hemoglobin %dA/A changes, we opted not to regress the hemodynamic signal out of the fluorescence signal.

### Event detection and quantification in neuronal fluorescence data

Optical detection of cortical hemodynamics and calcium dynamics has been extensively cross-validated as a surrogate for routine electrophysiology for detecting CSD, offering rich spatial information about traveling waves.^29–32^. The temporal bounds of seizure and CSD events were identified by taking the derivative of the root mean square of the neuronal fluorescence signal at each pixel. We then used the MATLAB function ‘findchangepts’ to find abrupt signal changes at the start and end of seizure and CSD events. We used an event detection threshold of %dF/F signal rising greater than 20% of baseline. Note, typical widefield %dF/F (fluorescence change) values are +/- 5-10% during routine physiological fluctuations in a mouse. In contrast, seizure and CSD are such large events that %dF/F fluorescence typically rises by 50-200%. The event detection threshold of 20% was determined through trial and error to accurately identify the bounds of seizure and CSD. This approach was cross-validated against the current gold standard for electrographic CSD detection - visual scoring by a trained clinician^33^. Recordings (and pixels therein) not crossing this threshold were excluded from analysis. For each event (seizure or CSD) in each pixel, duration was calculated as the difference between start and end times, and amplitude was calculated as the peak %dF/F value between the start and end time. Then, average event amplitude was computed within right hemisphere, left hemisphere, and total brain space by averaging across pixels. Overall event durations were computed by comparing the event start times in the first and last pixel with right hemisphere, left hemisphere, and total brain space. For CSDs, the first pixel within the brain mask to have an event start time was used as a global reference point for computing time to peak at other pixels for spatial maps of CSD trajectory. For CSDs, the percentage of pixels from each hemisphere recruited into the event was calculated. Average within-hemisphere CSD areal propagation speed was computed by dividing the area recruited into the CSD by the event duration within each hemisphere. Average linear propagation speed was computed by identifying the minimum and maximum points within each hemisphere of the CSD trajectory maps and drawing a line between them. This line profile of the spatial gradient was fit with a linear regression from which the slope was computed and converted into mm/min. Of note, clinical seizure duration in human ECT does not consistently include the 0.5-8 second stimulation period, because EEG activity cannot be measured during electrical pulse delivery. Our mouse optical neuroimaging paradigm allows us to observe that high-amplitude cortical discharges occur during stimulus delivery; we thus included this activity as part of the seizure.

Both seizure and CSD are well known to trigger a period of suppressed brain activity. To calculate post-event suppression time, a short time Fourier transform (STFT) power spectrum was determined for each pixel and for average right/left hemisphere fluorescence. Hemispheric suppression endpoints were detected using the findchangepts function for the fluorescence signal in each hemisphere. The suppression end point for each pixel was computed by finding the inflection point of up-trending 1-3Hz power (associated with return of slow waves from anesthesia) that is nearest to the respective global suppression endpoint. Then, the suppression duration was computed by taking the difference between the suppression end time and the event (seizure or SD) end time. Suppression times were then averaged within each hemisphere.

The maximum intensity seizure pixel was identified for all end-titration (i.e., CSD- generating) seizures (n=42) by identifying the median value within the top 99% of pixel fluorescence amplitude in seizure maps. We then calculated the shortest distance between the seizure peak pixel and the nearest stimulation electrode (**Fig. 2i**), as well as the shortest distance between the seizure peak pixel and the CSD initiation reference pixel (**Fig. 2j**). Electrode-to-seizure and seizure-to-CSD distances were averaged across all 42 events. To assess the significance of these spatial relationships, a shuffle analysis was conducted by randomly permuting the data and recalculating the average distances for 10,000 iterations. The distribution of shuffled distances was compared to the actual distances, and statistically verified by t-test with Bonferroni correction for multiple comparisons.

### Statistical comparisons of seizure and CSD phenotypes

For analyses below summarizing all recorded seizure and CSD events, distribution normality testing was performed using the D’Agostino & Pearson test to determine use of parametric or non-parametric testing. We used non-parametric Kruskal-Wallis ANOVA with Dunn’s multiple comparison test for comparison of seizure laterality index between stimulation types (**Fig. 2c**), as well as comparison of seizure duration (**Extended Data Fig. 3a**) and amplitude (**Extended Data Fig. 3b**) across CSD outcomes. Post-event suppression times (**Extended Data Fig. 3c,d**) were compared using the parametric Brown-Forsythe ANOVA with Dunnett’s multiple comparison test. The relative contribution of the three types of stimulation configurations (right, left, bilateral) for right, left and bilateral CSDs were compared using Fisher’s exact test (**Fig. 2f**). Average CSD fluorescence amplitude (**Fig. 2g**) and duration (**Fig. 2h**) were compared using the non-parametric Mann-Whitney test.

### Mixed effects model of stimulation parameter titration

For analyses of the effect of titrating pulse parameters (frequency, current level) on seizure properties (overall duration, average mean fluorescence amplitude, **Fig. 3**), we excluded cases that did not cross the seizure threshold in any pixels, thus excluding all evoked responses at 1 mA. A mixed effects model was fit using the outcome of either seizure duration or average peak fluorescence amplitude (%dF/F), transformed on a log scale to achieve approximate normality. Frequency and current level were the independent variables. A model including an interaction between current level and frequency was fit and the interaction evaluated using an F-test with a Satterthwaite correction for the degrees of freedom. If the interaction was significant (p < 0.05) the model was refit for each current level and the effect of increasing frequency evaluated. Specific values of frequency were replaced with integer values (levels 1,2,3,4,5 correspond to 5, 10, 25, 50, 100 Hz) to succinctly evaluate trends in the outcome as a function of frequency. If the interaction was not significant, the procedure was repeated and the main effects of frequency and amplitude were evaluated marginally, followed by Wald tests of specific contrasts. Models of current and frequency effects included electrode placement, animal ID, and pulse counts as covariates to adjust for imbalances across conditions. Electrode placement and pulse count were tested in separate sensitivity analyses to preclude convergence problems with the mixed effects models due to the small number of animals. Similarly, frequency was included as an ordered variable in the model with pulse count.

### Time-to-event Model of CSD

To assess how modulating pulse parameters impacts the probability of CSD after seizure, we fit a Cox model substituting current level for time and including frequency as the independent variable and stratification by animal ID. Current is a surrogate for time in this model because an animal receives increasing current levels at each frequency until the event, CSD, is observed. Once a CSD was observed, no further increases in current were assessed. Frequency was considered both as an ordered variable, to assess the overall trend with increasing frequency, and as a categorical variable to describe the results at specific frequencies. Electrode placement is included in the models reported here and in sensitivity analyses substituting pulse count for electrode frequency. Hypothesis tests of variables with individual terms are based on Wald tests, and for those with multiple categories (electrode placement, frequency) on likelihood ratio tests (LRT). Hypothesis tests are two-sided with a Type I error rate of 0.05, uncorrected for multiple comparisons.

### Diffuse Optics and Vitals Monitoring in ECT Patients

All procedures described below were approved by the University of Pennsylvania Institutional Review Board for the observational cohort study, SWEET COMBO: Studying Waves Evoked by Electroconvulsive Therapy with Combined Optical Monitoring of Blood flow and Oxygenation. Patients included in the study (n=18) were recruited from the pool of patients already being treated with ECT at Pennsylvania Hospital (Penn Medicine Health System) between February and July 2024. All participants provided written informed consent to participate. Of note, participation in this cross-sectional study had no impact on which patients were treated with ECT, nor in what manner. All patients were treated with standard of care protocols for Pennsylvania Hospital using a Σigma device (SigmaStim). Study enrollees include 9 women and 9 men, ages ranging 23-82 (median age 60). Enrollees self-identified as white (n=14), black (n=3), Asian (n=1) and Hispanic (n=1).

During ECT treatment, in addition to routine clinical monitoring (EKG, EEG, vitals), two diffuse optical monitoring probes were placed on the forehead surface in the regions overlying right (F4) and left (F3) prefrontal cortex. The probes and optical system provide simultaneous measurement of blood flow index using diffuse correlation spectroscopy^34,35^ (DCS) and tissue oxygen saturation using frequency-domain diffuse optical spectroscopy^35,36^ (FD-DOS) from scalp, skull, and cortex underlying the probes (see **Supplemental Methods** on instrumentation)^35^. To improve accuracy of the recovered tissue oxygen saturation, FD-DOS combined data from four separations (1.5, 2, 2.5, and 3 cm) on the tissue surface to extract a single datapoint. For DCS flow monitoring, each optical probe utilized a long and short source-detector pair that were fit independently. The light from the short separation pair (1 cm) penetrates scalp and skull, while light from the long separation pair (2.5 cm) probes deeper to the cortex. The optical property data collected concurrently with FD-DOS was input to the DCS fitting to account for changes in absorption and scattering that may otherwise confound the recovered blood flow index. Briefly, FD-DOS and DCS data were fit using the semi-infinite, homogenous solutions to the frequency domain-photon diffusion equation and the correlation diffusion equation, respectively. The interested reader is directed to reference^34^ for greater detail on the theoretical models used. In the main text, we only show blood flow obtained from the long separation pair that is sensitive to cortex (example short separation data is shown in **Extended Data Fig. 5** and **Supplemental Discussion**).

Diffuse optical recordings started from a pre-stimulation baseline period and lasted for several minutes after treatment; the treatment period had variable duration depending on the procedure room schedule, effects of anesthesia, and patient response. Routine intermittent vitals monitoring (collected at 2-minute intervals) was supplemented with continuous vitals monitoring (20 second intervals) using a noninvasive finger monitor for cardiac output (Acumen IQ cuff; HemoSphere monitor, Edwards)^37^. A total of n=27 recordings were performed on n=17 patients **(Extended Data Table 2** and **3**). Fifteen of these recordings were excluded because the recording signal did not meet quality control standards (see **Supplemental Discussion/Methods**). The remaining n=12 recordings are presented in **Fig. 4** and **Extended Data Fig. 4**. Based on prior measurements in cross-validated CSDs^38–42^, we defined CSD as a post-ictal wave of at least 2-minute duration before return to 100% of baseline, and with peak >200% of baseline blood flow and/or with >±5% fluctuation around baseline brain oxygen saturation. For recordings where bilateral sensor data was available, if these criteria were met in both hemispheres, the CSDs were classified as bilateral; if criteria were met in only one hemisphere, CSD was classified as unilateral.

**Fig. 4.**
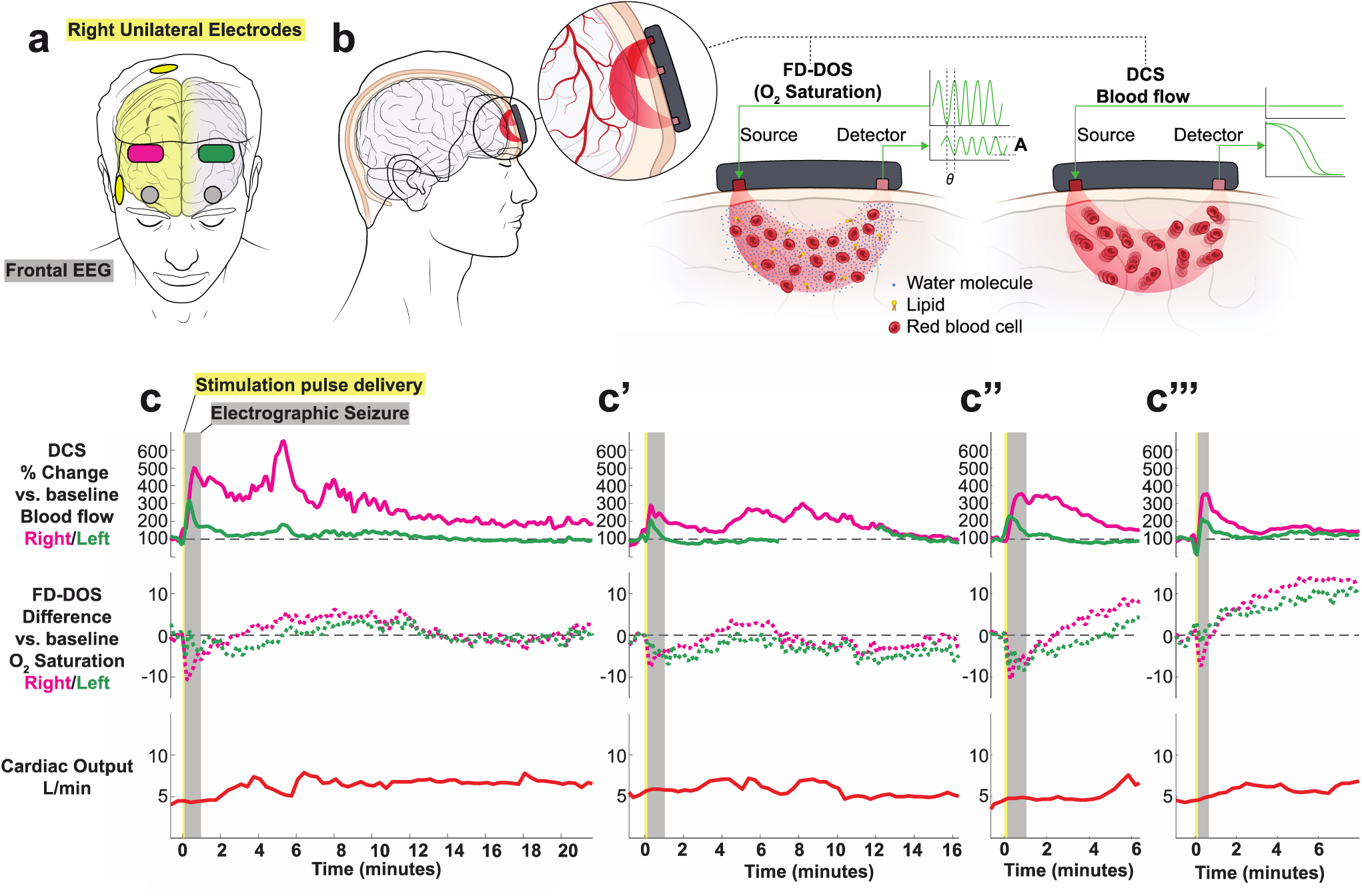
Cerebral hemodynamics during four ECT sessions from a single patient exhibit expected features of post-ictal CSD. **a**, Diagram of diffuse optical probes (magenta on right at F4, green on left at F3), frontal EEG (grey circles at FP1 and FP2), and right unilateral electrodes (yellow). **b**, Simplified schematic of non-invasive diffuse optics sensors, using paired red-light sources and detectors to regionally measure both deep (brain) and superficial (skin, scalp) blood flow (diffuse correlation spectroscopy, DCS) from two source-detector separations of 1 cm and 2.5 cm and oxygen saturation (frequency-domain diffuse optical spectroscopy, FD-DOS) from four source-detector separations of 1.5, 2, 2.5, and 3 cm. For clarity, the co-located DCS and FD-DOS probes are separated in the schematic. See **Methods** and **Supplemental Discussion and Methods** on diffuse optics. **c,** Series of four recordings on four different treatment days from the same subject receiving right unilateral ECT. Note, the blood flow data shown are derived from the source-detector pair with largest separation and thus largest depth penetration into the brain (Data at shorter separations are given in **Extended Data Fig. 5**). The oxygen saturation data is the result of combining data from all FD-DOS source-detector pairs for improved signal quality.

## Results

### ECT seizure is followed by cortical spreading depolarization in mice

We created a new mouse ECT model to facilitate optical neuroimaging of large-scale brain activity during ECT stimulation. We used transgenic *Thy1-jRGECO1a* mice which express the red fluorescent calcium sensor jRGECO1a in excitatory neurons^43^. Mice were implanted with a transparent polymer window overlying the intact skull to enable optical access to activity-dependent dynamics across the cortical surface. We measured neuronal activity with jRGECO1a fluorescence as well as hemodynamic activity with optical intrinsic signal (OIS) imaging, using green light illumination and a 1-photon widefield fluorescence mesoscope (optical set-up shown in **Fig. 1b**, spectra shown in **Extended Data Fig. 1**). To facilitate concurrent electrical stimulation within the imaging field, each mouse cranial window was implanted with five brass electrodes stereotactically affixed to the skull surface overlying frontal, temporo-parietal, and occipital cortices (**Fig. 1a**). Brain activity was recorded with widefield imaging under dexmedetomidine anesthesia during 153 ECT stimulation blocks. To broadly survey the stimulation parameter space (see Methods and **Extended Data Table 1**), we controlled the stimulation pulse parameters using an isolated pulse stimulator, with spatial configuration varied by changing which pair of electrodes were stimulated. For each set of tested conditions, stimulation was titrated by delivering successive rounds of pulses with increasing current (1, 2, 5, 10, 25 mA, **Fig. 1c**) followed by a monitoring period before restimulating.

Representative neuronal calcium dynamics during ECT titration are shown in **Fig. 1**. Low amplitude current steps (1 and 2 mA) elicited a small <15% dF/F evoked response and then an immediate return of slow wave activity characteristic of anesthesia (**Fig. 1c**). At higher current (5, 10 mA), ECT elicited seizure activity (high-amplitude, irregular discharges generalized across the cortex), followed by 5-10 seconds of post-ictal suppression, and then return of slow wave activity. At 25 mA, seizure was followed by a qualitatively distinct electrical event: cortical spreading depolarization (CSD, see **Extended Data Video 1**, depicted as still frames in **Fig. 1c** and time series in **Fig. 1c”**). CSD is a slow-traveling, high-amplitude wave of electrochemical depolarization that can be detected with high spatiotemporal fidelity using all-optical detection of neural dynamics or hemodynamics^29–32^. Post-ictal CSD was observed as a slowly propagating, high-amplitude wavefront of neuronal calcium elevation that concentrically expanded across the whole brain over the course of ∼160 seconds, followed by a prolonged period of suppressed neuronal calcium dynamics. All 38/38 ECT titration sessions reached the threshold to trigger post-ictal CSD. Direct current electrocorticography (DC-ECoG) in a different rodent ECT model likewise demonstrates post-ictal CSD in a current-thresholded manner (see **Extended Data Fig. 2, Supplemental Discussion**).

Note, despite their magnitude, CSDs are virtually invisible on routine alternating current (AC)-amplified EEG, as they are physically obscured by volume conduction and spatial blurring through the scalp and skull, and are digitally hidden with high-pass filtering (>0.5 Hz)^33,44,45^. Notably, subjecting mouse brain fluorescence data to the same >0.5 Hz high-pass filtering rendered CSD waves invisible on widefield imaging, and at point measurements throughout the brain appeared as a flat line that is otherwise indistinguishable from post-ictal suppression (see **Extended Data Video 1**).

### Electrode configuration shapes the spatial topography of seizure and CSD

We then considered whether systematically varying electrode configuration would impact the spatial topography of both seizure and post-ictal CSD waves. In our mouse model with five electrodes, ten electrode pairs were possible for left unilateral (L, green), right unilateral (R, magenta), or bilateral (BL, black) stimulation. Intuitively, we observed that electrode configuration shaped the spatial topography of both the evoked seizure (**Fig. 2a**, mapping seizure fluorescence amplitude at each pixel) as well as the trajectory of the subsequent CSD traveling wave (**Fig. 2b**, mapping time-to-peak of CSD at each pixel). Bilateral electrode placement produced relatively symmetric seizures, while R or L unilateral electrode placement recruited asymmetric seizure with higher amplitude in the stimulated hemisphere than the contralateral hemisphere (**Fig. 2c,d**, L vs R p**** < 0.0001, L vs BL p**** < 0.0001, R vs BL p** =0.0026).

In contrast to preceding seizures, post-ictal CSDs occurred in an all-or-none fashion anatomically, recruiting either ∼0% or ∼100% of cortical pixels in a given hemisphere (**Fig. 2b,e**). Trials that evoked bilateral CSD waves were disproportionately triggered by bilateral stimulation, while trials that evoked unilateral CSD waves were disproportionately triggered by same-side unilateral electrodes (**Fig. 2f**, p < 0.0001). Bilateral and unilateral CSDs tended to achieve similar fluorescence amplitude (**Fig. 2g,** p= 0.331), but bilateral CSD events tended to be longer in duration (**Fig. 2h,** p = 0.0011), primarily because of cases of asynchronous onset of multifocal CSD between hemispheres (e.g., **Fig. 2b** frontotemporal configuration 1,2 in panel 1, where CSD in L hemisphere peaked ∼1 minute before the R hemisphere). CSD propagation was noted to originate variably from both singleton and multiple foci, with all foci inevitably expanding to invade the entire hemisphere in which they started, but never traveling across midline. On average within any given hemisphere, we observed that CSD wavefronts spread at 4.1 mm/min (± standard deviation of 0.9 mm/min), consistent with prior electrophysiology. CSDs expanded in 2D at an average rate of 13.2 mm^2^/minute (± standard deviation of 3.6 mm^2^/min). Seizure, while always generalized bilaterally throughout cortex, tended to exhibit highest amplitude at a short distance from the stimulation electrodes (**Fig. 2a,i**, p**** < 0.0001), consistent with prior predictions from brain electric field models for various stimulation geometries^5^. Likewise, CSD initial foci tended to occur in close spatial proximity to pixels exhibiting peak fluorescence values during the preceding seizure (**Fig. 2b,j**, p**** < 0.0001).

### Pulse parameters modulate seizure and post-ictal CSD

Next, we examined titration data to test how pulse parameters, current (mA) and frequency (Hz), influence seizure and CSD. For each treatment session, individual mice were assigned to a pulse frequency and spatial configuration, and then serially stimulated with increasing current steps, monitoring the evoked seizures at each step with optical imaging, and ending at the current threshold where seizure led to CSD. For each seizure, we measured the maximum amplitude (peak fluorescence intensity averaged across all brain pixels, **Fig. 3a**) and overall event duration across the brain (**Fig. 3b**). Current titrations in individual mice at a fixed frequency and electrode configuration are represented as individual lines in **Fig. 3a,b**, with titration curves ending at the current threshold where the evoked seizure triggered a CSD. Some combinations of frequency and current were not tested (e.g., the combination of 100 Hz stimulation at 25 mA was not reached in any titration because all mice experienced CSD at lower current steps). Trials that did not meet our amplitude threshold for defining a seizure (see Methods) were excluded from statistical analysis. To assess how pulse current and frequency modify brain outcomes, we fit a mixed effects model (see Methods), using a random term to account for repeated measures from each animal and fixed effects for the assigned electrode configuration and pulse count.

For every mouse titration, across all tested frequencies, seizure amplitude went up with increasing current (note positive slope to each titration line shown in **Fig. 3a**). The effect of current differed by frequency (p< 0.001 F-test, with LRT interaction vs. main effects model). Increasing pulse frequency significantly increased the amplitude of the resultant seizure, producing steeper slopes in titrations for higher frequency conditions of 50 and 100 Hz shown in **Fig. 3a**. Within individual current levels, frequency did not significantly modulate seizure amplitude at low current steps (2 mA, p=0.63), but each level increase in pulse frequency increased seizure amplitude by a factor of 1.2 fold, significantly so at 5 mA (p< 0.001) and 10 mA (p=0.001), but not at 25 mA (p=0.11), see **Fig. 3a**. Electrode placement did not significantly influence seizure amplitude, either in the interaction model (p=0.70) or the individual models (p>0.05). In sensitivity analyses, higher pulse count did not significantly contribute to current and frequency effects (p=0.45 LRT). Higher frequency pulses thus elicit higher amplitude seizures independent of other variables.

In the same seizure titrations, increasing current led to increased seizure duration (**Fig. 3b,** p< 0.001), in a manner that did not vary significantly by pulse frequency condition (p=0.39). Compared to seizures at the 2 mA current step, seizure duration increased 2.8- fold at 5 mA (p< 0.001), 3.5-fold at 10 mA (p< 0.001) and 2.8-fold at 25 mA (p=0.005) when controlling for individual frequency level and electrode placement. Somewhat surprisingly, though increasing frequency led to larger seizure amplitude, seizures became briefer. Compared to 5 Hz stimulation, seizure duration decreased by a factor of 0.6 for 10 Hz (p=0.08), by 0.4 at 10 Hz (p=0.004), and by a factor of 0.3 at 50 and 100 Hz (p< 0.001). Seizures generally increased in duration with increasing current, but seizures at CSD threshold often had reduced duration, apparently because the expanding CSD terminated the ictal state. Frequency (p< 0.001) and pulse count (p=0.006) were associated with longer seizure duration. Delivery of 50 pulses yielded 1.9-fold longer seizure duration than 25 pulses (p=0.006) and 100 pulses yielded 2.4 fold longer seizure duration than 25 pulses (p=0.012). Duration at pulse counts of 50 and 100 did not differ significantly from each other. (p=0.49)

These parameters also impact the probability of generating CSD. Seizures that were sufficient to elicit CSD had significantly higher seizure amplitudes **(Extended Data Fig. 3**, p < 0.0001 for both unilateral and bilateral CSD), but no difference in seizure duration compared to seizures that do not elicit CSD. We thus asked if higher frequency stimulation (which elicits higher amplitude seizures) was more like to result in a CSD. To this end, we built a Cox model with current level as an analog of time (sequentially increased from 1, 2, 5, 10, 25 mA until CSD is achieved) and adjusting for electrode placement and pulse count. We find that increasing the stimulation frequency by one level (e.g., 5 to 10 Hz) increased the cumulative probability of CSD 2.1-fold at any point in the titration (95% CI 1.6-2.7, p< 0.001, **Fig. 3c**). Neither electrode placement (p=0.199) nor pulse count (p=0.64) was associated with the probability of CSD in this model. Compared to the lowest intensity 5 Hz stimulation, the probability of CSD was 16.5-fold greater at 100 Hz (p < 0.001) and 3.5-fold greater, but not significantly so, at 50 Hz (p=0.23); 10 Hz (p=0.87) and 25 Hz (p=0.18) did not differ from 5 Hz. Thus, at any current level, higher stimulation frequency statistically increased the likelihood of eliciting CSD, effectively leading CSD to occur earlier in a titration at lower current. Lastly, we quantified post-event suppression time by measuring the time between the terminal event (seizure or CSD) and subsequent return of baseline slow wave 1-3 Hz spectral power from anesthesia (**Extended Data Fig. 3c,d**). We observed that post-ictal suppression times were relatively brief (20-70 seconds before return of slow waves), but suppression times were significantly longer in hemispheres affected by CSD (110-270s; see Figure for pairwise comparisons.)

Using concurrent hemodynamic imaging, we observed that seizure elicited a blush of hyperemia. During CSD, this blush was followed by subsequent traveling waves of hypo- and hyperperfusion, which propagated slowly in the wake of the neuronal depolarization wave, consistent with prior reports^31,46^. Intuitively, we observe that bilateral CSD consistently triggers bilateral hemodynamic traveling waves (**Extended Data Video 2**), unilateral CSD triggers hemodynamic waves in only one hemisphere (**Extended Data Video 3**), and seizure without CSD generates the initial blush, but no post-ictal traveling hemodynamic wave (**Extended Data Video 4**). In each of these videos, two ROI pixel time series are presented to illustrate that CSD hemodynamic waves can occur with a variable post-ictal time delays and morphology, depending on where in the brain one takes a point measurement in the path of a multifocal traveling wave. Optical detection of this hemodynamic wave is a reliable biomarker of CSD in both rodents and humans^29,46–50^.

### Cerebral hemodynamics in human ECT patients show expected features of post-ictal CSD

Given that CSDs were reliably induced in mice across a wide range of stimulation parameters, we hypothesized that CSDs could also occur in human patients receiving ECT. We further hypothesized that the hemodynamic waves of post-ictal CSD could be translationally detected using non-invasive diffuse optical monitoring of cerebral blood flow and oxygen saturation (**Fig. 4a)**. To this end, we measured local cerebral hemodynamics using multimodal optical sensors on the forehead during routine human ECT treatments in a single-center observational cohort study. We define four criteria for detecting CSD in these measurements. First, we expect to see an initial hemodynamic wave during seizure; subsequently during the post-ictal period, we expect hemodynamics will either return to baseline in cases/hemispheres without CSD, or exhibit a second wave (consistent with Extended Data Videos 1, 3, 4 and prior reports^38^). Second, based on prior invasive regional measurements in human patients, if this post-ictal wave is a CSD, it will exhibit >2-minute long duration of 200-600% increase in cerebral blood flow relative to baseline^39^. Third, based on prior oximetry during CSD, we expect post-ictal waves will exhibit variable polyphasic waves of either hypoxic, hyperoxic, or mixed response,^40,41^ with amplitude modulation of ±5% from baseline^42^. Because flow data was not available in all subjects, we consider meeting either criterion two (blood flow) or three (oximetry) sufficient. Fourth, given the impact of electrode spatial configuration on the hemispheric laterality of mouse CSDs (**Fig. 2**), we predict that right unilateral ECT will elicit post-ictal hemodynamic waves that are primarily right-sided (and occasionally bilateral), while bilateral stimulation will elicit primarily bilateral waves.

In this pilot study, we obtained n=12 recordings of bilateral cerebral hemodynamics during right unilateral ECT (n=5 patients) and bitemporal ECT (n=5 patients), summarized in **Extended Data Table 2** and **3**. We first present recordings from four treatments on different days from an index course of right unilateral ECT for a 50-year-old woman with treatment-resistant depression (**Fig. 4b)**. All four treatments resulted in ∼60s, bilateral seizures on frontal EEG (grey shaded region). During seizure, relative cerebral blood flow (measured with DCS long source-detector separation) increased by 200-500%. This change was observed predominantly in the right hemisphere, with an associated 5-10% *decrease* in brain %O_2_ saturation (derived from the FD-DOS signal).

After the seizure ended electrographically (i.e., scored by a clinician as a flat line on EEG), all four recordings exhibited an inflection point and then a distinct post-ictal wave of 200- 600% increase in relative blood flow that lasted for over 2-minutes. Interestingly, unlike the preceding seizure, this second hyperemic wave was associated with a 5-10% *increase* in brain oxygen saturation, suggesting a hyperoxic surge of blood flow that kept pace with the metabolic demand of the brain^40,47^. These regional measurements of post- ictal hemodynamic waves exhibited variable waveform morphology and time delays relative to the initial electrographic seizure. In the first three recordings, post-ictal hyperemic waves were restricted to the right hemisphere (magenta lines), and in the last recording the hemodynamic changes occurred bilaterally, indicating three right hemispheric post-ictal events and one bilateral event. These hemispheric comparisons provide a within-subject control to demonstrate that hemodynamic changes are due to local brain metabolism and not to global autonomic changes. To further corroborate that these 200-600% changes in blood flow index are not due to systemic circulation, we implemented non-invasive cardiac output monitoring throughout the procedure. We observe that systemic blood flow from the heart was relatively constant at 5 ± 2 L/min throughout the procedure. Taken together, these waves meet all four of our criteria for defining post-ictal CSD. Of note, over this interval, the patient exhibited a relatively typical clinical response to ECT. At her pre-treatment evaluation she reported intermittent suicidal ideation, anhedonia, excess sleep, feelings of worthlessness, hopelessness, and low energy. After 2 weeks (6 treatments) of ECT, she reported her mood, energy, and sleep had improved, and that her suicidal thoughts had resolved. She denied any side effects of treatment.

We collected eight more recordings from n=8 patients (**Extended Data Fig. 4**) and similarly observed post-ictal hyperemic waves meeting criteria for CSD in an additional 3/3 recordings of right unilateral ECT as well as 4/5 recordings of bitemporal ECT. In all recordings where sensor data was available bilaterally, putative CSDs occurred bilaterally. This post-ictal surge in cerebral blood flow was also evident in the superficial short separation DCS detectors, with variable amplitude relative to the long separation DCS detectors (see **Extended Data Fig. 5** and **Supplemental Discussion**). Finally, we provide additional replication cases obtained independently at another institution using a commercial continuous-wave functional near infrared spectroscopy (fNIRS) instrument to measure cerebral oximetry in human ECT patients (see **Extended Data Fig. 6** and **Supplemental Discussion**). Thus, the hemodynamic signatures of post-ictal CSD were reliably observed in three different modalities of non-invasive optical monitoring.

## Discussion

In this study, we show that electroconvulsive therapy (ECT) reliably elicits post-ictal cortical spreading depolarization (CSD) in both mouse models and in human patients. This discovery unifies several previously disconnected observations: (1) that ECT induces lasting inhibitory plasticity in brain activity^7^, (2) that therapeutic response to ECT requires high energy seizures of sufficient magnitude^51^, and (3) that stimulation parameters modulate seizure and clinical outcomes from ECT^20^. Our findings suggest that strong stimulation elicits seizure of sufficient magnitude to generate post-ictal CSD.

Spreading depolarization may engage mechanisms of inhibitory plasticity that contribute to the brain’s clinical response to ECT. Post-ictal CSD has primarily been observed with intracranial electrocorticography (ECoG) in both human patients^52–54^ and in animal models, dating back to Leão’s original demonstration that CSD can be induced by direct electrical or mechanical stimulation of cortex^55–57^. Because of the requirement for invasive neurosurgery for electrophysiologic detection of CSD, virtually all reports of post-ictal CSD in human patients have been obtained in individuals with brain lesions that already required surgery (e.g., stroke and traumatic brain injury). In the context of injury and metabolically fragile brain tissue, CSD is known to exacerbate excitotoxicity and worsen clinical outcomes. However, in the context of seizure in an uninjured brain, preclinical evidence suggests that CSD may act as an intrinsic protective mechanism, terminating the seizure and inducing inhibitory plasticity that prevents future seizures^38,58^. CSD is not known to be harmful in brains without underlying injury; in this context, CSD has been observed to paradoxically induce growth and plasticity genes, downregulate cell death genes, and protect against subsequent injury^54,59–62^. It remains unknown whether ECT-induced CSDs are harmful or potentially part of the therapeutic mechanism. This cannot be answered with this pilot study’s limited sample of 12 recordings over single treatments (nearly all of which exhibited evidence of CSD). This will be tested in future large-scale clinical studies. One potential clue is the observation that post-ictal blood flow waves were associated with a 5-10% *increase* in cerebral O_2_ saturation – a feature associated with CSDs with preserved neurovascular coupling and adequate perfusion, in contrast to the *decreased* O_2_ saturation seen in CSDs from ischemic injuries^47^. If CSD proves to be part of the therapeutic mechanism of ECT, this would invite the exciting possibility of bypassing seizure to directly trigger CSD. Indeed, we find that certain stimulation conditions (i.e., brief trains of high frequency pulses) more efficiently generate CSD with minimal seizure duration, while low frequency pulses require higher levels of current to elicit high-amplitude seizure or CSD.

The inhibitory, antiseizure effects of CSD might help explain the long-known inhibitory effects of ECT. ECT raises the seizure threshold in over 90% of patients^7^, requiring clinicians to progressively increase stimulation intensity between treatments to maintain adequate seizure^11^. Ironically, this inhibitory effect makes ECT highly effective at treating status epilepticus, particularly in super-refractory cases when maximal anti-seizure pharmacotherapy has failed^63–65^. Furthermore, in those treated with ECT, EEG demonstrates sustained increases in aperiodic power spectral density (an index of cortical inhibitory tone)^66,67^, as well as suppressed evoked measures of excitability (transcranial magnetic stimulation evoked potentials)^10^. Human PET imaging has likewise shown that ECT increases cortical GABA concentrations^68^, and rodent studies have demonstrated corresponding changes in GABA release^69^ and GABA receptor mRNA levels^70^. In the immediate aftermath of CSD, synaptic activity is potently suppressed, initially by depolarization block of axonal action potentials, then by inhibition of presynaptic neurotransmitter release due to activation of adenosine A1 receptors^71,72^. Adenosine accumulation is also observed with seizures^73^ and ECT^74^. Within inhibitory circuits, ECT-induced plasticity may require cell-type specific molecular regulators of circuit excitability^75^ and act on excitability of single neurons^76^. We hypothesize that post-ictal CSD may contribute to these multi-scale forms of inhibitory plasticity after ECT.

To our knowledge, this is the first study to record brain activity during ECT pulse delivery. We used optical recordings to circumvent electrical signal artifacts that confound routine EEG monitoring during stimulation. Previous computational predictions have shown that larger evoked electric fields correlate with improved clinical response^15,77,78^, which we speculate may be due to increased probability of a large electric field triggering CSD. Real-time recording of brain activity during ECT may prove valuable for future studies on stimulus parameter optimization, just as it has for transcranial magnetic stimulation (TMS)^79–81^, deep brain stimulation (DBS)^82,83^, and ultrasound neuromodulation^84^. Of note, in our mouse model, we observed that high frequency stimulation at 100 Hz sometimes evoked only a single high-amplitude spike during pulse delivery, without persistent post-stimulation ictal activity. We opted to include evoked activity during stimulation as part of the seizure, though the physiology governing persistent post-stimulation seizure may differ from acute evoked discharges during stimulation. Both brief high-amplitude discharges during stimulation as well as persistent post-stimulation seizure were sufficient to generate CSD in our mouse model.

Post-ictal CSD has the potential to explain many epiphenomena seen in other measurements of brain activity from ECT patients. We observed that larger seizures were more likely to generate CSD in mice, which could theoretically explain why higher EEG ictal spectral power is associated with improved clinical outcomes in humans^12,14,15^. Our rodent data also showed longer post-event suppression time after CSD compared to seizure alone, which could explain why longer suppression times are associated with improved patient outcomes^12,85^ and longer latency to reorientation^86^. Extended EEG recording after treatment may be necessary to validate if post-ictal suppression is associated with CSD^45,87^. Looking beyond EEG, CSD is known to produce lasting changes in brain hemodynamics, which could account for why effective ECT seizures elicit lasting perfusion changes 1-hour after treatment^88^ as well as on longer timescales^89,90^. Furthermore, we observed that right unilateral stimulation primarily generated unilateral post-ictal waves. Unilateral CSD could play a role in ECT-induced volumetric changes that occur preferentially in the stimulated hemisphere and correlate with clinical outcomes and brain network functional connectivity changes^91–96^. CSDs are so named because neocortex is most accessible for recording brain activity. Indeed, this is a key limitation of our optical methods for recording cortical dynamics - spreading depolarization can also occur in subcortical structures, which might have important implications for both the clinical benefits and side effects of ECT.

We speculate that post-ictal CSD has been occurring undetected during the millions of treatments that have been performed over the last 86 years. In clinical practice, EEG is typically turned off after the seizure terminates, and furthermore, EEG cannot reliably detect CSDs ^33,44^ unless the skull has been penetrated^45,97^. This is thought to be due to digital high-pass filtering of slow waveforms, as well as spatial smearing from volume conduction of CSD dynamics through the brain, skull, and scalp. Very focal CSDs with narrow-width wavefronts may be particularly challenging to detect^45^. Optical recordings in rodent models have long proven a valuable surrogate for CSD detection, providing richer spatial detail on traveling waves compared to electrographic recordings^29,46–50^. Here we replicate these findings in both optical imaging of neural and blood dynamics, and provide supplemental verification of CSD with DC-ECoG (Extended Data Figure 2). However, non-invasive optical methods have not yet been extended to human subjects prior to this work.

Indeed, Dreier et al. have proposed that non-invasive optical hemodynamic measures should prove sufficient to detect CSD in human patients^97^. Here we find that optical features of CSD can be detected non-invasively at patient bedside, and meet all of our criteria for defining CSD based on prior literature on the amplitude, duration, and morphology of CSD hemodynamic waves^38–42^. This approach to CSD detection may have valuable clinical applications in neurocritical care for individuals without intracranial ECoG implants. Previous studies have characterized the hemodynamic features of CSD using invasive probes in the brain^39–41,98^ in those with brain injuries. In migraine patients, CSD has been detected non-invasively using functional MRI^42,99^ and radiotracer/PET imaging^100,101^, revealing that CSD is the physiologic basis of migraine aura (and may perhaps explain headache as an occasional side effect of ECT). Note, PET and MRI are incompatible with ECT due to the need for mechanical ventilation during treatment. In this study we implemented a non-invasive investigational device that combined frequency-domain diffuse optical spectroscopy (FD-DOS) to measure tissue oxygenation with diffuse correlation spectroscopy (DCS) to capture blood flow rate changes.

Our human optical recordings have several limitations. First, it is not possible to obtain electrographic verification of post-ictal CSD in these recordings – which would require invasive neurosurgery for ECoG implantation, and would not be clinically appropriate in ECT patients. As expected from prior reports^33,44,45,97^, we were unable to detect CSD using scalp DC-EEG (see **Supplement**). Second, our approach relied on regional point measurements over non-hair bearing skin. Future investigations using broader spatial coverage at multiple points would be needed to detect CSD propagation across a brain hemisphere. Third, signal quality is significantly influenced by patient motion, skin temperature, and variation in skull and skin microanatomy that can influence the detected optical path distribution (see **Supplemental Discussion**/**Methods**). These factors can introduce systematic errors that influence absolute tissue optical properties. We have ameliorated these effects by computing relative change in blood flow and oxygenation indices within each individual.

This study implemented a new mouse model of ECT that enabled experimental control over stimulation combined with real-time monitoring of brain activity with widefield imaging. To model the three standard ECT electrode placements (right unilateral, bitemporal, bifrontal), we attached five electrodes to the intact skull with silver-conductive epoxy to test ten configurations of electrode pairs. This builds on prior innovation using screw electrodes in rats, which have been shown to have superior translational validity to conventional rodent ECT with auricular (ear clip) electrodes.^102^ Skull electrodes help control for the confounding effects of current shunting through the low-resistance skin^103^. Indeed, we observed the use of ear clip electrodes required 5-10 times more current to elicit seizure or CSD compared to skull electrodes (see **Supplement**). In addition, we substituted a conventional rodent ECT device for a broadly customizable isolated pulse stimulator, enabling the first systematic evaluation of ECT pulse parameters in an animal model. We found that increasing pulse frequency increased seizure amplitude and probability of CSD, while low frequency pulses elicited long duration seizures without increased probability of CSD. This builds on prior studies showing that high frequency electrical pulses can elicit CSD in hippocampal slices^104^, and that low intensity stimulation does not elicit CSD^105^. One limitation of this study is its inability to exhaustively survey the virtually infinite space of pulse parameters and electrode configurations^20^. We opted to survey multiple electrode and pulse parameters simultaneously, using pseudorandomization to sample stimulation conditions in a balanced fashion, and linear mixed effects modeling to isolate pulse parameter-specific effects in titration data. This is an imperfect solution, and future investigations would benefit from systematically modulating individual stimulation parameters, as well as testing for specific effects in translational models of neuropsychiatric disease. Lastly, using a current titration strategy, we identified that increasing pulse frequency lowered the current threshold required to elicit a seizure of a given magnitude. This has important ramifications for novel ECT protocols that modulate current amplitude^26,78,106^. Future investigations should explore whether high frequency stimulation is a more current-sparing approach of achieving therapeutic efficacy. This study thus provides a translational framework for measuring brain activity to optimize ECT stimulation parameters and precisely control seizure and post-ictal CSD. These findings further highlight opportunities to modernize ECT and directly target treatment-induced mechanisms of brain plasticity.

## Supporting information

Manuscript Main Text

Supplemental Methods and Discussion

## Author Contributions

All authors helped with data interpretation and editorial feedback on the paper.

**ZPR** conceptualized overall project; designed, performed, and analyzed results of mouse experiments; generated Penn IRB protocol, collected and analyzed human data; wrote original manuscript, produced figures; contributed funding.

**JBM** contributed to diffuse optics method development, human data collection and analysis, and text in the original manuscript

**AS** contributed to Penn mouse imaging data analysis and text in the original manuscript

**DKQ** designed and performed UNM human recordings

**BEL** contributed UNM mouse recordings and text in the original manuscript.

**MEP** conceptualized and implemented time-to-event and linear mixed effects modeling and contributed text in the original manuscript.

**AK** contributed to the statistical analysis presented in the original manuscript.

**CGF** contributed to methodological development of human studies and IRB protocol.

**WBB** contributed to conceptualization and methods development for diffuse optics.

**GH** is an ECT clinician who helped supervise human data collection.

**JPR** is an ECT clinician who helped supervise human data collection.

**MAC** is an ECT clinician who helped supervise human data collection and advise on project conceptualization.

**YIS** helped supervise development of human studies and funding acquisition.

**CWS** conceptualized and supervised UNM mouse experiments.

**CCA** is an ECT clinician and conceptualized and supervised UNM human case series.

**AGY** contributed diffuse optics resources for human recordings, method development, data analysis, postdoctoral supervision and funding support of JBM.

**EMG** contributed to method development of mouse ECT model and widefield imaging, editorial supervision of original manuscript, postdoctoral supervision of ZPR.

## Acknowledgements

We would like to thank the patients who participated in the study for their generosity and openness. This work was supported by the Penn Psychiatry Residency Research Track award NIH R25MH119043 (ZPR); Institute for Translational Medicine and Therapeutics of the University of Pennsylvania NIH UL1TR001878 (ZPR); NIH P41-EB029460 (AGY); NIH NS078805, NS051288, NS070680 (BEL); NIH NS106901 (CWS); NIH P50HD105354 (MEP, AK). The authors would like to thank Emily Hoddeson for assistance with mouse husbandry and genotyping, Dr. Adam Bauer for advice on mouse optical methods, Dr. Jin-Moo Lee for sharing breeders from the *Thy1-jRGECO1a* mouse line, ANT Neuro for loaning DC-EEG equipment, Katy Brown for facilitating human continuous vitals monitoring, Dr. Holly Lisanby for advice on project conceptualization, Jurre Blom for assistance with manuscript illustrations, Dr. Brian White for help launching the collaboration between the diffuse optics and ECT research teams, and Dr. Rodrigo M. Forti for help with data interpretation and diffuse optical instrumentation.

## Competing Interests

No financial conflicts or completing interests to disclose.

## Data Availability

Source Data are provided in the Goldberg Lab gnode repository (https://gin.g-node.org/GoldbergNeuroLab).

## Notes

https://gin.g-node.org/GoldbergNeuroLab/

### Competing Interest Statement

The authors have declared no competing interest.

### Summary of Updates

Revised text for improved clarity, additional analyses suggested by reviewers

## References

1. Espinoza, R. T. & Kellner, C. H. Electroconvulsive Therapy. N Engl J Med 386, 667–672, (2022).

2. Kaster, T. S., Blumberger, D. M., Gomes, T., Sutradhar, R., Wijeysundera, D. N. & Vigod, S. N. Risk of suicide death following electroconvulsive therapy treatment for depression: a propensity score-weighted, retrospective cohort study in Canada. Lancet Psychiatry 9, 435–446, (2022).

3. Fink, M. What was learned: studies by the consortium for research in ECT (CORE) 1997-2011. Acta Psychiatr Scand 129, 417–426, (2014).

4. Sackeim, H. A. Modern Electroconvulsive Therapy: Vastly Improved yet Greatly Underused. JAMA Psychiatry 74, 779–780, (2017).

5. Deng, Z. D., Robins, P. L., Regenold, W., Rohde, P., Dannhauer, M. & Lisanby, S. H. How electroconvulsive therapy works in the treatment of depression: is it the seizure, the electricity, or both? Neuropsychopharmacology 49, 150–162, (2024).

6. Regenold, W. T., Noorani, R. J., Piez, D. & Patel, P. Nonconvulsive Electrotherapy for Treatment Resistant Unipolar and Bipolar Major Depressive Disorder: A Proof-of-concept Trial. Brain Stimul 8, 855–861, (2015).

7. Duthie, A. C., Perrin, J. S., Bennett, D. M., Currie, J. & Reid, I. C. Anticonvulsant Mechanisms of Electroconvulsive Therapy and Relation to Therapeutic Efficacy. J ECT 31, 173–178, (2015).

8. Lloyd, R. L. & Sattin, A. The Behavioral Physiology and Antidepressant Mechanisms of Electroconvulsive Shock. J ECT 31, 159–166, (2015).

9. Voineskos, D., Levinson, A. J., Sun, Y., Barr, M. S., Farzan, F., Rajji, T. K., Fitzgerald, P. B., Blumberger, D. M. & Daskalakis, Z. J. The Relationship Between Cortical Inhibition and Electroconvulsive Therapy in the Treatment of Major Depressive Disorder. Sci Rep 6, 37461, (2016).

10. Bajbouj, M., Lang, U. E., Niehaus, L., Hellen, F. E., Heuser, I. & Neu, P. Effects of right unilateral electroconvulsive therapy on motor cortical excitability in depressive patients. J Psychiatr Res 40, 322–327, (2006).

11. Krystal, A. D., Coffey, C. E., Weiner, R. D. & Holsinger, T. Changes in seizure threshold over the course of electroconvulsive therapy affect therapeutic response and are detected by ictal EEG ratings. J Neuropsychiatry Clin Neurosci 10, 178–186, (1998).

12. Perera, T. D., Luber, B., Nobler, M. S., Prudic, J., Anderson, C. & Sackeim, H. A. Seizure expression during electroconvulsive therapy: relationships with clinical outcome and cognitive side effects. Neuropsychopharmacology 29, 813–825, (2004).

13. Zhang, J. Y., Xu, S. X., Zeng, L., Chen, L. C., Li, J., Jiang, Z. Y., Tan, B. J., Gu, C. L., Lai, W. T., Kong, X. M., Wang, J., Rong, H. & Xie, X. H. Improved Safety of Hybrid Electroconvulsive Therapy Compared With Standard Electroconvulsive Therapy in Patients With Major Depressive Disorder: A Randomized, Double-Blind, Parallel-Group Pilot Trial. Front Psychiatry 13, 896018, (2022).

14. Miller, J., Jones, T., Upston, J., Deng, Z. D., McClintock, S. M., Ryman, S., Quinn, D. & Abbott, C. C. Ictal Theta Power as an Electroconvulsive Therapy Safety Biomarker: A Pilot Study. J ECT 38, 88–94, (2022).

15. Miller, J., Jones, T., Upston, J., Deng, Z. D., McClintock, S. M., Erhardt, E., Farrar, D. & Abbott, C. C. Electric Field, Ictal Theta Power, and Clinical Outcomes in Electroconvulsive Therapy. Biol Psychiatry Cogn Neurosci Neuroimaging 8, 760–767, (2023).

16. Gillving, C., Ekman, C. J., Hammar, A., Landen, M., Lundberg, J., Movahed Rad, P., Nordanskog, P., von Knorring, L. & Nordenskjold, A. Seizure Duration and Electroconvulsive Therapy in Major Depressive Disorder. JAMA Netw Open 7, e2422738, (2024).

17. Khadka, N., Deng, Z. D., Lisanby, S. H., Bikson, M. & Camprodon, J. A. Computational Models of High-Definition Electroconvulsive Therapy for Focal or Multitargeting Treatment. J ECT, (2024).

18. Argyelan, M. et al. Electroconvulsive therapy-induced volumetric brain changes converge on a common causal circuit in depression. Mol Psychiatry 29, 229–237, (2024).

19. Deng, Z. D., Argyelan, M., Miller, J., Quinn, D. K., Lloyd, M., Jones, T. R., Upston, J., Erhardt, E., McClintock, S. M. & Abbott, C. C. Electroconvulsive therapy, electric field, neuroplasticity, and clinical outcomes. Mol Psychiatry 27, 1676–1682, (2022).

20. Peterchev, A. V., Rosa, M. A., Deng, Z. D., Prudic, J. & Lisanby, S. H. Electroconvulsive therapy stimulus parameters: rethinking dosage. J ECT 26, 159–174, (2010).

21. Sackeim, H. A., Prudic, J., Nobler, M. S., Fitzsimons, L., Lisanby, S. H., Payne, N., Berman, R. M., Brakemeier, E. L., Perera, T. & Devanand, D. P. Effects of pulse width and electrode placement on the efficacy and cognitive effects of electroconvulsive therapy. Brain Stimul 1, 71–83, (2008).

22. Wang, X., Padawer-Curry, J. A., Bice, A. R., Kim, B., Rosenthal, Z. P., Lee, J.-M., Goyal, M. S., Macauley, S. L. & Bauer, A. Q. Spatiotemporal relationships between neuronal, metabolic, and hemodynamic signals in the awake and anesthetized mouse brain. Cell Reports 43, (2024).

23. Brier, L. M., Landsness, E. C., Snyder, A. Z., Wright, P. W., Baxter, G. A., Bauer, A. Q., Lee, J. M. & Culver, J. P. Separability of calcium slow waves and functional connectivity during wake, sleep, and anesthesia. Neurophotonics 6, 035002, (2019).

24. Mitra, A., Kraft, A., Wright, P., Acland, B., Snyder, A. Z., Rosenthal, Z., Czerniewski, L., Bauer, A., Snyder, L., Culver, J., Lee, J. M. & Raichle, M. E. Spontaneous Infra-slow Brain Activity Has Unique Spatiotemporal Dynamics and Laminar Structure. Neuron 98, 297–305 e296, (2018).

25. Mukhtar, F., Regenold, W. & Lisanby, S. H. Recent advances in electroconvulsive therapy in clinical practice and research. Faculty Reviews 12, (2023).

26. Peterchev, A. V., Krystal, A. D., Rosa, M. A. & Lisanby, S. H. Individualized Low-Amplitude Seizure Therapy: Minimizing Current for Electroconvulsive Therapy and Magnetic Seizure Therapy. Neuropsychopharmacology 40, 2076–2084, (2015).

27. Brier, L. M. & Culver, J. P. Open-source statistical and data processing tools for wide-field optical imaging data in mice. Neurophotonics 10, 016601, (2023).

28. Franklin, K. B. J. & Paxinos, G. The Mouse Brain in Stereotactic Coordinates. (Academic Press, 2012).

29. Chung, D. Y., Sugimoto, K., Fischer, P., Bohm, M., Takizawa, T., Sadeghian, H., Morais, A., Harriott, A., Oka, F., Qin, T., Henninger, N., Yaseen, M. A., Sakadzic, S. & Ayata, C. Real-time non-invasive in vivo visible light detection of cortical spreading depolarizations in mice. J Neurosci Methods 309, 143–146, (2018).

30. Balbi, M., Vanni, M. P., Silasi, G., Sekino, Y., Bolanos, L., LeDue, J. M. & Murphy, T. H. Targeted ischemic stroke induction and mesoscopic imaging assessment of blood flow and ischemic depolarization in awake mice. Neurophotonics 4, 035001, (2017).

31. Zhao, H. T., Tuohy, M. C., Chow, D., Kozberg, M. G., Kim, S. H., Shaik, M. A. & Hillman, E. M. C. Neurovascular dynamics of repeated cortical spreading depolarizations after acute brain injury. Cell Rep 37, 109794, (2021).

32. Smith, S. E. et al. Astrocyte deletion of alpha2-Na/K ATPase triggers episodic motor paralysis in mice via a metabolic pathway. Nat Commun 11, 6164, (2020).

33. Hofmeijer, J., van Kaam, C. R., van de Werff, B., Vermeer, S. E., Tjepkema-Cloostermans, M. C. & van Putten, M. Detecting Cortical Spreading Depolarization with Full Band Scalp Electroencephalography: An Illusion? Front Neurol 9, 17, (2018).

34. Carp, S. A., Robinson, M. B. & Franceschini, M. A. Diffuse correlation spectroscopy: current status and future outlook. Neurophotonics 10, 013509, (2023).

35. Durduran, T., Choe, R., Baker, W. B. & Yodh, A. G. Diffuse Optics for Tissue Monitoring and Tomography. Rep Prog Phys 73, (2010).

36. Zhou, X., Xia, Y., Uchitel, J., Collins-Jones, L., Yang, S., Loureiro, R., Cooper, R. J. & Zhao, H. Review of recent advances in frequency-domain near-infrared spectroscopy technologies [Invited]. Biomed Opt Express 14, 3234–3258, (2023).

37. Kouz, K., et al. Intraoperative hypotension when using hypotension prediction index software during major noncardiac surgery: a European multicentre prospective observational registry (EU HYPROTECT). BJA Open 6, 100140, (2023).

38. Tamim, I., Chung, D. Y., de Morais, A. L., Loonen, I. C. M., Qin, T., Misra, A., Schlunk, F., Endres, M., Schiff, S. J. & Ayata, C. Spreading depression as an innate antiseizure mechanism. Nat Commun 12, 2206, (2021).

39. Lemale, C. L., Luckl, J., Horst, V., Reiffurth, C., Major, S., Hecht, N., Woitzik, J. & Dreier, J. P. Migraine Aura, Transient Ischemic Attacks, Stroke, and Dying of the Brain Share the Same Key Pathophysiological Process in Neurons Driven by Gibbs-Donnan Forces, Namely Spreading Depolarization. Front Cell Neurosci 16, 837650, (2022).

40. Winkler, M. K., Dengler, N., Hecht, N., Hartings, J. A., Kang, E. J., Major, S., Martus, P., Vajkoczy, P., Woitzik, J. & Dreier, J. P. Oxygen availability and spreading depolarizations provide complementary prognostic information in neuromonitoring of aneurysmal subarachnoid hemorrhage patients. J Cereb Blood Flow Metab 37, 1841–1856, (2017).

41. Hecht, N., Haddad, D., Neumann, K., Schumm, L., Dengler, N. F., Wessels, L., Domer, P., Helgers, S., Meinert, F., Major, S., Lemale, C. L., Dreier, J. P., Vajkoczy, P. & Woitzik, J. Reduced brain oxygen response to spreading depolarization predicts worse outcome in ischaemic stroke. Brain, (2024).

42. Hadjikhani, N., Sanchez Del Rio, M., Wu, O., Schwartz, D., Bakker, D., Fischl, B., Kwong, K. K., Cutrer, F. M., Rosen, B. R., Tootell, R. B., Sorensen, A. G. & Moskowitz, M. A. Mechanisms of migraine aura revealed by functional MRI in human visual cortex. Proc Natl Acad Sci U S A 98, 4687–4692, (2001).

43. Dana, H., Novak, O., Guardado-Montesino, M., Fransen, J. W., Hu, A., Borghuis, B. G., Guo, C., Kim, D. S. & Svoboda, K. Thy1 transgenic mice expressing the red fluorescent calcium indicator jRGECO1a for neuronal population imaging in vivo. PLoS One 13, e0205444, (2018).

44. Dreier, J. P., Major, S., Lemale, C. L., Kola, V., Reiffurth, C., Schoknecht, K., Hecht, N., Hartings, J. A. & Woitzik, J. Correlates of Spreading Depolarization, Spreading Depression, and Negative Ultraslow Potential in Epidural Versus Subdural Electrocorticography. Front Neurosci 13, 373, (2019).

45. Chamanzar, A., George, S., Venkatesh, P., Chamanzar, M., Shutter, L., Elmer, J. & Grover, P. An Algorithm for Automated, Noninvasive Detection of Cortical Spreading Depolarizations Based on EEG Simulations. IEEE Trans Biomed Eng 66, 1115–1126, (2019).

46. Ayata, C., Shin, H. K., Salomone, S., Ozdemir-Gursoy, Y., Boas, D. A., Dunn, A. K. & Moskowitz, M. A. Pronounced hypoperfusion during spreading depression in mouse cortex. J Cereb Blood Flow Metab 24, 1172–1182, (2004).

47. Dreier, J. P. The role of spreading depression, spreading depolarization and spreading ischemia in neurological disease. Nat Med 17, 439–447, (2011).

48. de Crespigny, A., Rother, J., van Bruggen, N., Beaulieu, C. & Moseley, M. E. Magnetic resonance imaging assessment of cerebral hemodynamics during spreading depression in rats. J Cereb Blood Flow Metab 18, 1008–1017, (1998).

49. Zhou, C., Yu, G., Furuya, D., Greenberg, J., Yodh, A. & Durduran, T. Diffuse optical correlation tomography of cerebral blood flow during cortical spreading depression in rat brain. Opt Express 14, 1125–1144, (2006).

50. Yamato, H., Jin, T. & Nomura, Y. Near infrared imaging of intrinsic signals in cortical spreading depression observed through the intact scalp in hairless mice. Neurosci Lett 701, 213–217, (2019).

51. Sackeim, H. A., Prudic, J., Devanand, D. P., Kiersky, J. E., Fitzsimons, L., Moody, B. J., McElhiney, M. C., Coleman, E. A. & Settembrino, J. M. Effects of stimulus intensity and electrode placement on the efficacy and cognitive effects of electroconvulsive therapy. N Engl J Med 328, 839–846, (1993).

52. Fabricius, M., Fuhr, S., Willumsen, L., Dreier, J. P., Bhatia, R., Boutelle, M. G., Hartings, J. A., Bullock, R., Strong, A. J. & Lauritzen, M. Association of seizures with cortical spreading depression and peri-infarct depolarisations in the acutely injured human brain. Clin Neurophysiol 119, 1973–1984, (2008).

53. Dreier, J. P., Major, S., Pannek, H. W., Woitzik, J., Scheel, M., Wiesenthal, D., Martus, P., Winkler, M. K., Hartings, J. A., Fabricius, M., Speckmann, E. J., Gorji, A. & group, C. s. Spreading convulsions, spreading depolarization and epileptogenesis in human cerebral cortex. Brain 135, 259–275, (2012).

54. Dreier, J. P. et al. Recording, analysis, and interpretation of spreading depolarizations in neurointensive care: Review and recommendations of the COSBID research group. J Cereb Blood Flow Metab 37, 1595–1625, (2017).

55. Leao, A. A. Further observations on the spreading depression of activity in the cerebral cortex. J Neurophysiol 10, 409–414, (1947).

56. Leao, A. A. P. Spreading Depression of Activity in the Cerebral Cortex. Journal of Neurophysiology 7, 359–390, (1944).

57. Bragin, A., Penttonen, M. & Buzsaki, G. Termination of epileptic afterdischarge in the hippocampus. J Neurosci 17, 2567–2579, (1997).

58. Zakharov, A., Chernova, K., Burkhanova, G., Holmes, G. L. & Khazipov, R. Segregation of seizures and spreading depolarization across cortical layers. Epilepsia 60, 2386–2397, (2019).

59. Dell’Orco, M., Weisend, J. E., Perrone-Bizzozero, N. I., Carlson, A. P., Morton, R. A., Linsenbardt, D. N. & Shuttleworth, C. W. Repetitive spreading depolarization induces gene expression changes related to synaptic plasticity and neuroprotective pathways. Front Cell Neurosci 17, 1292661, (2023).

60. Nedergaard, M. & Hansen, A. J. Spreading depression is not associated with neuronal injury in the normal brain. Brain Res 449, 395–398, (1988).

61. Kobayashi, S., Harris, V. A. & Welsh, F. A. Spreading depression induces tolerance of cortical neurons to ischemia in rat brain. J Cereb Blood Flow Metab 15, 721–727, (1995).

62. Matsushima, K., Hogan, M. J. & Hakim, A. M. Cortical spreading depression protects against subsequent focal cerebral ischemia in rats. J Cereb Blood Flow Metab 16, 221–226, (1996).

63. Lambrecq, V., Villega, F., Marchal, C., Michel, V., Guehl, D., Rotge, J. Y. & Burbaud, P. Refractory status epilepticus: electroconvulsive therapy as a possible therapeutic strategy. Seizure 21, 661–664, (2012).

64. Garcia-Lopez, B. et al. Electroconvulsive Therapy in Super Refractory Status Epilepticus: Case Series with a Defined Protocol. Int J Environ Res Public Health 17, (2020).

65. Ahmed, J., Metrick, M., Gilbert, A., Glasson, A., Singh, R., Ambrous, W., Brown, L., Aykroyd, L. & Bobel, K. Electroconvulsive Therapy for Super Refractory Status Epilepticus. J ECT 34, e5–e9, (2018).

66. Smith, S. E., Ma, V., Gonzalez, C., Chapman, A., Printz, D., Voytek, B. & Soltani, M. Clinical EEG slowing induced by electroconvulsive therapy is better described by increased frontal aperiodic activity. Transl Psychiatry 13, 348, (2023).

67. Smith, S. E., Kosik, E. L., van Engen, Q., Kohn, J., Hill, A. T., Zomorrodi, R., Blumberger, D. M., Daskalakis, Z. J., Hadas, I. & Voytek, B. Magnetic seizure therapy and electroconvulsive therapy increase aperiodic activity. Transl Psychiatry 13, 347, (2023).

68. Sanacora, G., Mason, G. F., Rothman, D. L., Hyder, F., Ciarcia, J. J., Ostroff, R. B., Berman, R. M. & Krystal, J. H. Increased cortical GABA concentrations in depressed patients receiving ECT. Am J Psychiatry 160, 577–579, (2003).

69. Green, A. R. & Vincent, N. D. The effect of repeated electroconvulsive shock on GABA synthesis and release in regions of rat brain. Br J Pharmacol 92, 19–24, (1987).

70. Kang, I., Miller, L. G., Moises, J. & Bazan, N. G. GABAA receptor mRNAs are increased after electroconvulsive shock. Psychopharmacol Bull 27, 359–363, (1991).

71. Lindquist, B. E. & Shuttleworth, C. W. Adenosine receptor activation is responsible for prolonged depression of synaptic transmission after spreading depolarization in brain slices. Neuroscience 223, 365–376, (2012).

72. Lindquist, B. E. & Shuttleworth, C. W. Evidence that adenosine contributes to Leao’s spreading depression in vivo. J Cereb Blood Flow Metab 37, 1656–1669, (2017).

73. Boison, D. Adenosinergic signaling in epilepsy. Neuropharmacology 104, 131–139, (2016).

74. Lewin, E. & Bleck, V. Electroshock seizures in mice: effect on brain adenosine and its metabolites. Epilepsia 22, 577–581, (1981).

75. Chang, A. D. et al. Narp Mediates Antidepressant-Like Effects of Electroconvulsive Seizures. Neuropsychopharmacology 43, 1088–1098, (2018).

76. Ueta, Y., Yamamoto, R. & Kato, N. Layer-specific modulation of pyramidal cell excitability by electroconvulsive shock. Neurosci Lett 709, 134383, (2019).

77. Lee, W. H., Lisanby, S. H., Laine, A. F. & Peterchev, A. V. Comparison of electric field strength and spatial distribution of electroconvulsive therapy and magnetic seizure therapy in a realistic human head model. Eur Psychiatry 36, 55–64, (2016).

78. Abbott, C. C., Quinn, D., Miller, J., Ye, E., Iqbal, S., Lloyd, M., Jones, T. R., Upston, J., Deng, Z., Erhardt, E. & McClintock, S. M. Electroconvulsive Therapy Pulse Amplitude and Clinical Outcomes. Am J Geriatr Psychiatry 29, 166–178, (2021).

79. Gogulski, J., Ross, J. M., Talbot, A., Cline, C. C., Donati, F. L., Munot, S., Kim, N., Gibbs, C., Bastin, N., Yang, J., Minasi, C., Sarkar, M., Truong, J. & Keller, C. J. Personalized Repetitive Transcranial Magnetic Stimulation for Depression. Biol Psychiatry Cogn Neurosci Neuroimaging 8, 351–360, (2023).

80. Wang, J. B., Hassan, U., Bruss, J. E., Oya, H., Uitermarkt, B. D., Trapp, N. T., Gander, P. E., Howard, M. A., 3rd, Keller, C. J. & Boes, A. D. Effects of transcranial magnetic stimulation on the human brain recorded with intracranial electrocorticography. Mol Psychiatry 29, 1228–1240, (2024).

81. Peterchev, A. V., Goetz, S. M., Westin, G. G., Luber, B. & Lisanby, S. H. Pulse width dependence of motor threshold and input-output curve characterized with controllable pulse parameter transcranial magnetic stimulation. Clin Neurophysiol 124, 1364–1372, (2013).

82. Paulk, A. C., Zelmann, R., Crocker, B., Widge, A. S., Dougherty, D. D., Eskandar, E. N., Weisholtz, D. S., Richardson, R. M., Cosgrove, G. R., Williams, Z. M. & Cash, S. S. Local and distant cortical responses to single pulse intracranial stimulation in the human brain are differentially modulated by specific stimulation parameters. Brain Stimul 15, 491–508, (2022).

83. Huang, Y., Zelmann, R., Hadar, P., Dezha-Peralta, J., Richardson, R. M., Williams, Z. M., Cash, S. S., Keller, C. J. & Paulk, A. C. Theta-burst direct electrical stimulation remodels human brain networks. Nat Commun 15, 6982, (2024).

84. Murphy, K. R. et al. Optimized ultrasound neuromodulation for non-invasive control of behavior and physiology. Neuron, (2024).

85. Moulier, V., Guehl, J., Eveque-Mourroux, E., Quesada, P. & Rotharmel, M. A Retrospective Study of Postictal Suppression during Electroconvulsive Therapy. J Clin Med 11, (2022).

86. J, C. M. P., J, P. A. J. V., Stuiver, S., Aalbregt, E., Schmettow, M., Hofmeijer, J., van Waarde, J. A. & M, J. A. M. v. P. Seizure duration predicts postictal electroencephalographic recovery after electroconvulsive therapy-induced seizures. Clin Neurophysiol 148, 1–8, (2023).

87. Hickman, L. B., Kafashan, M., Labonte, A. K., Chan, C. W., Huels, E. R., Guay, C. S., Guan, M. J., Ching, S., Lenze, E. J., Farber, N. B., Avidan, M. S., Hogan, R. E. & Palanca, B. J. A. Postictal generalized electroencephalographic suppression following electroconvulsive therapy: Temporal characteristics and impact of anesthetic regimen. Clin Neurophysiol 132, 977–983, (2021).

88. Pottkamper, J. C. M., Verdijk, J., Aalbregt, E., Stuiver, S., van de Mortel, L., Norris, D. G., van Putten, M., Hofmeijer, J., van Wingen, G. A. & van Waarde, J. A. Changes in postictal cerebral perfusion are related to the duration of electroconvulsive therapy-induced seizures. Epilepsia 65, 177–189, (2024).

89. Hirano, J., Takamiya, A., Yamagata, B., Hotta, S., Miyasaka, Y., Pu, S., Iwanami, A., Uchida, H. & Mimura, M. Frontal and temporal cortical functional recovery after electroconvulsive therapy for depression: A longitudinal functional near-infrared spectroscopy study. J Psychiatr Res 91, 26–35, (2017).

90. Downey, D., Brigadoi, S., Trevithick, L., Elliott, R., Elwell, C., McAllister-Williams, R. H. & Anderson, I. M. Frontal haemodynamic responses in depression and the effect of electroconvulsive therapy. J Psychopharmacol 33, 1003–1014, (2019).

91. Mulders, P. C. R. et al. Structural changes induced by electroconvulsive therapy are associated with clinical outcome. Brain Stimul 13, 696–704, (2020).

92. Verdijk, J. et al. Longitudinal resting-state network connectivity changes in electroconvulsive therapy patients compared to healthy controls. Brain Stimul 17, 140–147, (2024).

93. Abbott, C. C., Jones, T., Lemke, N. T., Gallegos, P., McClintock, S. M., Mayer, A. R., Bustillo, J. & Calhoun, V. D. Hippocampal structural and functional changes associated with electroconvulsive therapy response. Transl Psychiatry 4, e483, (2014).

94. Argyelan, M. et al. Electric field causes volumetric changes in the human brain. Elife 8, (2019).

95. Cano, M., Lee, E., Cardoner, N., Martinez-Zalacain, I., Pujol, J., Makris, N., Henry, M., Via, E., Hernandez-Ribas, R., Contreras-Rodriguez, O., Menchon, J. M., Urretavizcaya, M., Soriano-Mas, C. & Camprodon, J. A. Brain Volumetric Correlates of Right Unilateral Versus Bitemporal Electroconvulsive Therapy for Treatment-Resistant Depression. J Neuropsychiatry Clin Neurosci 31, 152–158, (2019).

96. Leaver, A. M., Vasavada, M., Kubicki, A., Wade, B., Loureiro, J., Hellemann, G., Joshi, S. H., Woods, R. P., Espinoza, R. & Narr, K. L. Hippocampal subregions and networks linked with antidepressant response to electroconvulsive therapy. Mol Psychiatry 26, 4288–4299, (2021).

97. Drenckhahn, C., Winkler, M. K., Major, S., Scheel, M., Kang, E. J., Pinczolits, A., Grozea, C., Hartings, J. A., Woitzik, J., Dreier, J. P. & group, C. s. Correlates of spreading depolarization in human scalp electroencephalography. Brain 135, 853–868, (2012).

98. Thomas, R., Shin, S. S. & Balu, R. Applications of near-infrared spectroscopy in neurocritical care. Neurophotonics 10, 023522, (2023).

99. Gollion, C., Christensen, R. H., Ashina, H., Al-Khazali, H. M., Fisher, P. M., Amin, F. M., Lauritzen, M. & Ashina, M. Somatosensory migraine auras evoked by bihemispheric cortical spreading depression events in human parietal cortex. J Cereb Blood Flow Metab, 271678X241290606, (2024).

100. Woods, R. P., Iacoboni, M. & Mazziotta, J. C. Brief report: bilateral spreading cerebral hypoperfusion during spontaneous migraine headache. N Engl J Med 331, 1689–1692, (1994).

101. Olesen, J., Larsen, B. & Lauritzen, M. Focal hyperemia followed by spreading oligemia and impaired activation of rCBF in classic migraine. Ann Neurol 9, 344–352, (1981).

102. Theilmann, W., Loscher, W., Socala, K., Frieling, H., Bleich, S. & Brandt, C. A new method to model electroconvulsive therapy in rats with increased construct validity and enhanced translational value. J Psychiatr Res 53, 94–98, (2014).

103. Vöröslakos, M., Takeuchi, Y., Brinyiczki, K., Zombori, T., Oliva, A., Fernández-Ruiz, A., Kozák, G., Kincses, Z. T., Iványi, B., Buzsáki, G. & Berényi, A. Direct effects of transcranial electric stimulation on brain circuits in rats and humans. Nature Communications 9, (2018).

104. Su, Y., Radman, T., Vaynshteyn, J., Parra, L. C. & Bikson, M. Effects of high-frequency stimulation on epileptiform activity in vitro: ON/OFF control paradigm. Epilepsia 49, 1586–1593, (2008).

105. Turner, D. A., Degan, S., Galeffi, F., Schmidt, S. & Peterchev, A. V. Rapid, Dose-Dependent Enhancement of Cerebral Blood Flow by transcranial AC Stimulation in Mouse. Brain Stimul 14, 80–87, (2021).

106. Abbott, C. C. et al. Amplitude-determined seizure-threshold, electric field modeling, and electroconvulsive therapy antidepressant and cognitive outcomes. Neuropsychopharmacology 49, 640–648, (2024).

