## Supplementary material for "Electroconvulsive therapy generates a postictal wave of spreading depolarization in mice and humans": Manuscript Main Text

*See attached MP4 files.*

**Extended Data Video 1.** Comparison of full-band (unfiltered) versus 0.5 Hz high-pass filtered fluorescence dynamics during CSD. See corresponding still frames and time series in Figure 1. Time series represent data from an ROI pixel shown as a + on brain maps. Yellow shading indicates stimulation period, grey shading represents seizure persisting beyond the stimulation period. See methods text for pixel-wise seizure detection.

**Extended Data Video 2.** Comparison of neuronal fluorescence dynamics (top left) and hemodynamics (top right) from stimulation eliciting bilateral CSD. Same data as Extended Data Video 1. Time series represent data from ROI pixels shown as + and o on brain maps above.

**Extended Data Video 3.** Comparison of neuronal fluorescence dynamics (top left) and hemodynamics (top right) from stimulation eliciting unilateral CSD. Time series represent data from ROI pixels shown as + and o on brain maps above. Note high frequency stimulation (100 Hz) sometimes elicited a seizure so brief, it consisted of a single ictal discharge during the stimulation period, without persistent seizure after stimulation.

**Extended Data Video 4.** Comparison of neuronal fluorescence dynamics (top left) and hemodynamics (top right) from stimulation eliciting seizure alone, without CSD. Time series represent data from ROI pixels shown as + and o on brain maps above. Note total hemoglobin concentration returns to baseline within ~20 seconds after seizure alone in the absence of CSD.

| Mouse Number | Electrodes | Frequency (Hz) | Current step at CSD (mA) | Train Duration (s) | Pulse Count |
|---|---|---|---|---|---|
| 2 | 1,4 | 5 | 10 | 10 | 50 |
| 3 | 4,5 | 5 | 10 | 10 | 50 |
| 4 | 1,4 | 5 | 25 | 5 | 25 |
| 4 | 1,3 | 5 | 25 | 10 | 50 |
| 5 | 3,4 | 5 | 5 | 10 | 50 |
| 6 | 1,5 | 5 | 10 | 10 | 50 |
| 8 | 3,5 | 5 | 25 | 10 | 50 |
| 9 | 4,5 | 5 | 25 | 5 | 25 |
| 9 | 1,5 | 5 | 25 | 10 | 50 |
| 5 | 4,5 | 10 | 25 | 2.5 | 25 |
| 7 | 4,5 | 10 | 25 | 5 | 50 |
| 8 | 1,4 | 10 | 10 | 2.5 | 25 |
| 10 | 1,4 | 10 | 25 | 5 | 50 |
| 1 | 4,5 | 25 | 25 | 2 | 50 |
| 6 | 4,5 | 25 | 25 | 1 | 25 |
| 6 | 1,4 | 25 | 25 | 2 | 50 |
| 4 | 4,5 | 50 | 10 | 1 | 50 |
| 7 | 1,4 | 50 | 10 | 0.5 | 25 |
| 9 | 1,4 | 50 | 5 | 1 | 50 |
| 10 | 4,5 | 50 | 25 | 0.5 | 25 |
| 1 | 2,3 | 100 | 5 | 1 | 100 |
| 1 | 1,4 | 100 | 10 | 0.25 | 25 |
| 2 | 1,3 | 100 | 5 | 1 | 100 |
| 3 | 1,4 | 100 | 5 | 1 | 100 |
| 4 | 1,5 | 100 | 5 | 1 | 100 |
| 4 | 2,4 | 100 | 10 | 0.5 | 50 |
| 5 | 1,4 | 100 | 5 | 0.5 | 50 |
| 5 | 1,2 | 100 | 5 | 1 | 100 |
| 5 | 1,5 | 100 | 10 | 0.5 | 50 |
| 6 | 2,5 | 100 | 5 | 1 | 100 |
| 7 | 2,4 | 100 | 5 | 1 | 100 |
| 8 | 3,5 | 100 | 5 | 1 | 100 |
| 8 | 1,2 | 100 | 5 | 0.5 | 50 |
| 8 | 4,5 | 100 | 10 | 0.5 | 50 |
| 9 | 1,3 | 100 | 5 | 0.5 | 50 |
| 9 | 4,5 | 100 | 10 | 1 | 100 |
| 10 | 3,5 | 100 | 5 | 0.5 | 50 |
| 10 | 3,4 | 100 | 5 | 1 | 100 |

**Extended Data Table 1.** Stimulation parameters for Mouse ECT titration recordings.

| Patient | Stimulation | DOS Signal Quality | DCS Signal Quality | CSD wave (Left) | CSD wave (Right) | QC notes | Treatment Type / # | | Pulse Width (ms) | Frequency (Hz) | Pulse train duration (s) | Current (mA) | Impedance (Ohms) | Energy (J) | Motor seizure (s) | EEG/ Central Seizure (s) |
|---|---|---|---|---|---|---|---|---|---|---|---|---|---|---|---|---|
| 1 | BT | poor | poor | | | Poor sensor contact with skin | Index | 3 | 0.5 | 50 | 6 | 800 | 154 | 28.9 | 34 | 37 |
| 1 | BT | poor | poor | | | Poor sensor contact with skin | Index | 4 | 0.5 | 80 | 6 | 800 | 154 | 45.7 | 28 | 32 |
| 2 | RUL | good | poor | YES | YES | DCS censored for beta fluctuation | Index | 4 | 0.37 | 120 | 8 | 800 | 184 | 82.8 | 17 | 41 |
| 2 | RUL | poor | poor | | | Poor signal - beta fluctuation | Index | 5 | 0.37 | 120 | 8 | 800 | 177 | 80.9 | 11 | 41 |
| 3 | BT | N/A | N/A | | | File saving error | Maintenance | 19 | 0.5 | 90 | 8 | 800 | 132 | 61.3 | 41 | 61 |
| 4 | BF | poor | poor | | | Midline placement with BF electrodes | Maintenance | 29 | 0.5 | 120 | 6 | 800 | 167 | 77.4 | 54 | 79 |
| 4 | BT | poor | poor | | | DCS censored for beta fluctuation | Maintenance | 38 | 0.5 | 120 | 6 | 800 | 134 | 61.9 | 55 | 73 |
| 6 | BT | good | good | YES | YES | | Maintenance | 33 | 0.5 | 90 | 8 | 800 | 192 | 88.1 | 43 | 68 |
| 8 | RUL | good (R) | good (R) | - | YES | L sensor placed on back of head, too | Maintenance | 18 | 0.37 | 120 | 8 | 800 | 197 | 89.4 | 46 | 60 |
| 9 | RUL | poor | poor | | | Incompatible skull anatomy, low photon | Index | 5 | 0.37 | 120 | 8 | 800 | 162 | 58.4 | 22 | 76 |
| 9 | RUL | poor | poor | | | Incompatible skull anatomy, low photon | Index | 6 | 0.37 | 120 | 8 | 800 | 147 | 65.6 | 35 | 62 |
| 9 | RUL | poor | poor | | | Incompatible skull anatomy, low photon | Index | 7 | 0.37 | 120 | 8 | 800 | 176 | 78.7 | 28 | 51 |
| 11 | BT | good (R) | good (R) | YES | YES | | Maintenance | 38 | 0.5 | 120 | 6 | 800 | 203 | 92 | 46 | 69 |
| 12 | BT | good | poor | NO | NO | DCS censored for beta fluctuation | Maintenance | 10 | 0.5 | 90 | 8 | 800 | 125 | 58 | 40 | 65 |
| 12 | BT | poor | poor | | | DCS censored for beta fluctuation | Maintenance | 11 | 0.5 | 90 | 8 | 800 | 151 | 70.7 | 37 | 79 |
| 13 | RUL | poor | poor | | | Confounding motion artifact from | Index | 1 | 0.37 | 30 | 2 | 800 | 240 | 5.2 | 3 | 5 |
| 13 | RUL | good | good | NO | YES | | Index | 2 | 0.37 | 60 | 6 | 800 | 218 | 28.9 | 36 | 59 |
| 13 | RUL | good | good | NO | YES | | Index | 3 | 0.37 | 120 | 8 | 800 | 240 | 107.4 | 29 | 60 |
| 13 | RUL | good | good | NO | YES | | Index | 4 | 0.37 | 120 | 8 | 800 | 230 | 101.8 | 35 | 54 |
| 13 | RUL | good | good | YES | YES | | Index | 5 | 0.37 | 120 | 8 | 800 | 225 | 101 | 12 | 36 |
| 14 | BT | good | good | YES | YES | | Maintenance | 57 | 0.5 | 80 | 5 | 800 | 176 | 45.1 | 84 | 104 |
| 15 | RUL | good | good | YES | YES | | Maintenance | 9 | 0.37 | 120 | 8 | 800 | 237 | 105.7 | 44 | 57 |
| 16 | RUL | poor | poor | | | DCS censored for beta fluctuation | Index | 1 | 0.3 | 30 | 2 | 800 | 215 | 4.8 | 64 | 137 |
| 16 | RUL | poor | poor | | | Poor signal - beta fluctuation | Index | 2 | 0.3 | 50 | 5 | 800 | 187 | 17.1 | 81 | 105 |
| 16 | RUL | poor | poor | | | Poor signal - beta fluctuation | Index | 3 | 0.3 | 60 | 6 | 800 | 191 | 25.7 | 49 | 85 |
| 17 | BT | good | good | YES | YES | | Maintenance | 21 | 0.5 | 90 | 5 | 800 | 150 | 42.7 | 101 | 138 |
| 18 | RUL | poor | poor | | | Poor signal - beta fluctuation | Maintenance | 49 | 0.37 | 120 | 8 | 800 | 250 | 111.7 | 24 | 46 |

**Extended Data Table 2.** Stimulation parameters for Human ECT recordings.

| Patient | Stimulation | DOS Signal Quality | DCS Signal Quality | CSD wave (Left) | CSD wave (Right) | Methohexital (pre) | Methohexital (post) | Labetalol (pre) | Labetalol (post) | Ketorolac | Rocuronium | Succinylcholine | Glycopyrrolate | Dexmedetomidine | Sugammadex | Remifentanyl | Ondansetron | Propofol | Acetaminophen |
|---|---|---|---|---|---|---|---|---|---|---|---|---|---|---|---|---|---|---|---|
| 1 | BT | poor | poor | | | 200 | | 20 | | 30 | 80 | | | | 800 | | | | |
| 1 | BT | poor | poor | | | 200 | | 20 | | 30 | 80 | | | | 800 | | | | |
| 2 | RUL | good | poor | YES | YES | 100 | | 15 | | 30 | | 80 | 0.1 | | | | | | |
| 2 | RUL | poor | poor | | | 100 | | 15 | | 30 | | 80 | 0.1 | | | | | | |
| 3 | BT | N/A | N/A | | | 40 | | 15 | | | | 30 | | | | | | | |
| 4 | BF | poor | poor | | | 90 | | | | 15 | | 100 | | | | | | | |
| 4 | BT | poor | poor | | | 90 | | | | 15 | | 100 | | | | | | | |
| 6 | BT | good | good | YES | YES | 60 | | 15 | | | | 60 | | | | 100 | | | 650 |
| 8 | RUL | good (R) | good (R) | - | YES | 30 | | 35 | 15 | | | 50 | | | | 100 | | 30 | |
| 9 | RUL | poor | poor | | | 190 | | | | 30 | | 180 | | | | | | | |
| 9 | RUL | poor | poor | | | 190 | | | | 30 | | 180 | | | | | | | |
| 9 | RUL | poor | poor | | | 190 | | | | 30 | | 180 | | | | | | | |
| 11 | BT | good (R) | good (R) | YES | YES | 80 | | | | | | 70 | | | | | | | |
| 12 | BT | good | poor | NO | NO | 40 | | | | 30 | | 90 | | | | 150 | | | |
| 12 | BT | poor | poor | | | 40 | | | | 30 | | 90 | | | | 150 | | | |
| 13 | RUL | poor | poor | | | 180 | | 20 | | | | 100 | 0.1 | | | | | | |
| 13 | RUL | good | good | NO | YES | 120 | | 10 | | | | 100 | 0.2 | | | | | | |
| 13 | RUL | good | good | NO | YES | 120 | 20 | 15 | | | | 100 | 0.2 | | | | | | |
| 13 | RUL | good | good | NO | YES | 120 | 20 | 15 | | | | 100 | 0.2 | | | | | | |
| 13 | RUL | good | good | YES | YES | 120 | 20 | 15 | | | | 100 | 0.2 | | | | | | |
| 14 | BT | good | good | YES | YES | 90 | | 10 | | 30 | | 50 | | 20 | | 50 | | | |
| 15 | RUL | good | good | YES | YES | 90 | | 25 | 25 | | | 60 | | | | | 4 | | |
| 16 | RUL | poor | poor | | | 150 | | | | 30 | | 100 | | | | | | | |
| 16 | RUL | poor | poor | | | 100 | 20 | 25 | | | | 80 | | | | | | | |
| 16 | RUL | poor | poor | | | 140 | | 25 | | | | 80 | | | | | | | |
| 17 | BT | good | good | YES | YES | 40 | | | | | | 60 | | 10 | | 75 | | | |
| 18 | RUL | poor | poor | | | 50 | | | | | | 70 | | | | 75 | | | |

**Extended Data Table 3.** Medications administered for Human ECT recordings (milligram units for all medications except remifentanil and dexmedetomidine, which are in micrograms).

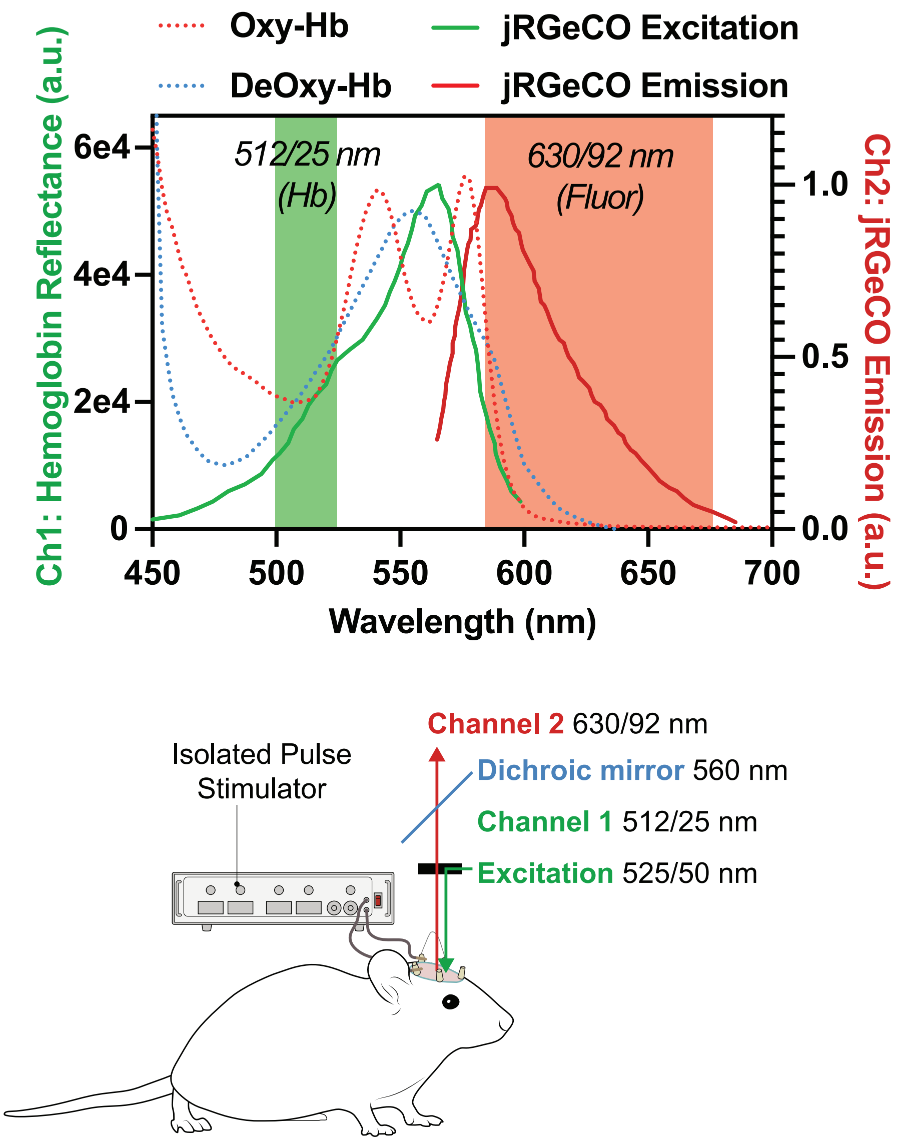


**Extended Data Figure 1**. Absorption/Reflectance spectra of oxy- and deoxy-hemoglobin, as well as excitation and emission spectra for jRGECO1a fluorescence. Illumination for both hemoglobin and fluorescence signals is provided at 525/50nm; light reflected and emitted by the brain is then directed through a dichroic and image splitting optics to isolate channel 1, green 512/25nm reflectance at the isosbestic point of oxy- and deoxy-hemoglobin (total blood volume/hemoglobin independent of oxygen saturation), and channel 2, red 630/92 nm fluorescence from jRGECO.

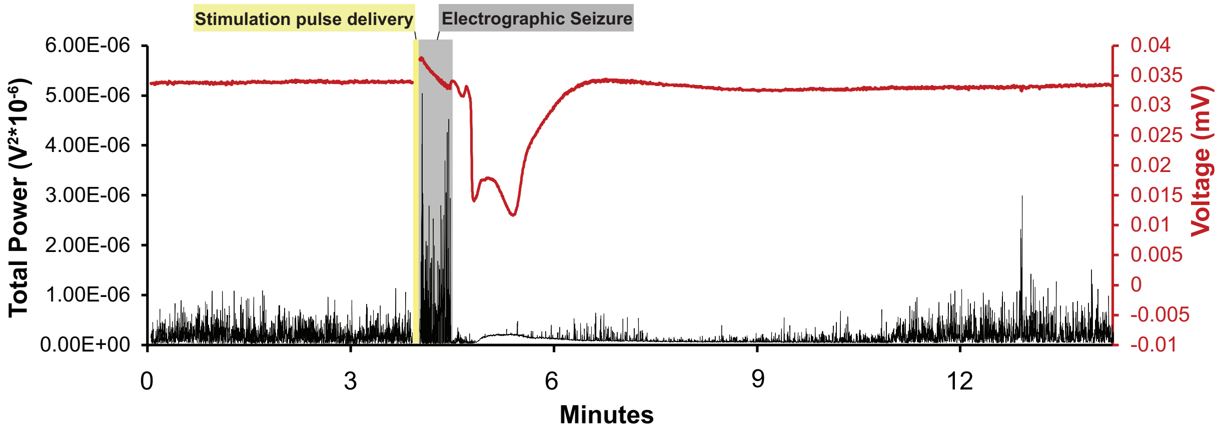


**Extended Data Figure 2.** Mouse Electrocorticography during ECT-induced seizure and CSD. Black trace corresponds to total power (0 - 500 Hz). Red trace corresponds to 2 Hz low-pass filtered data.

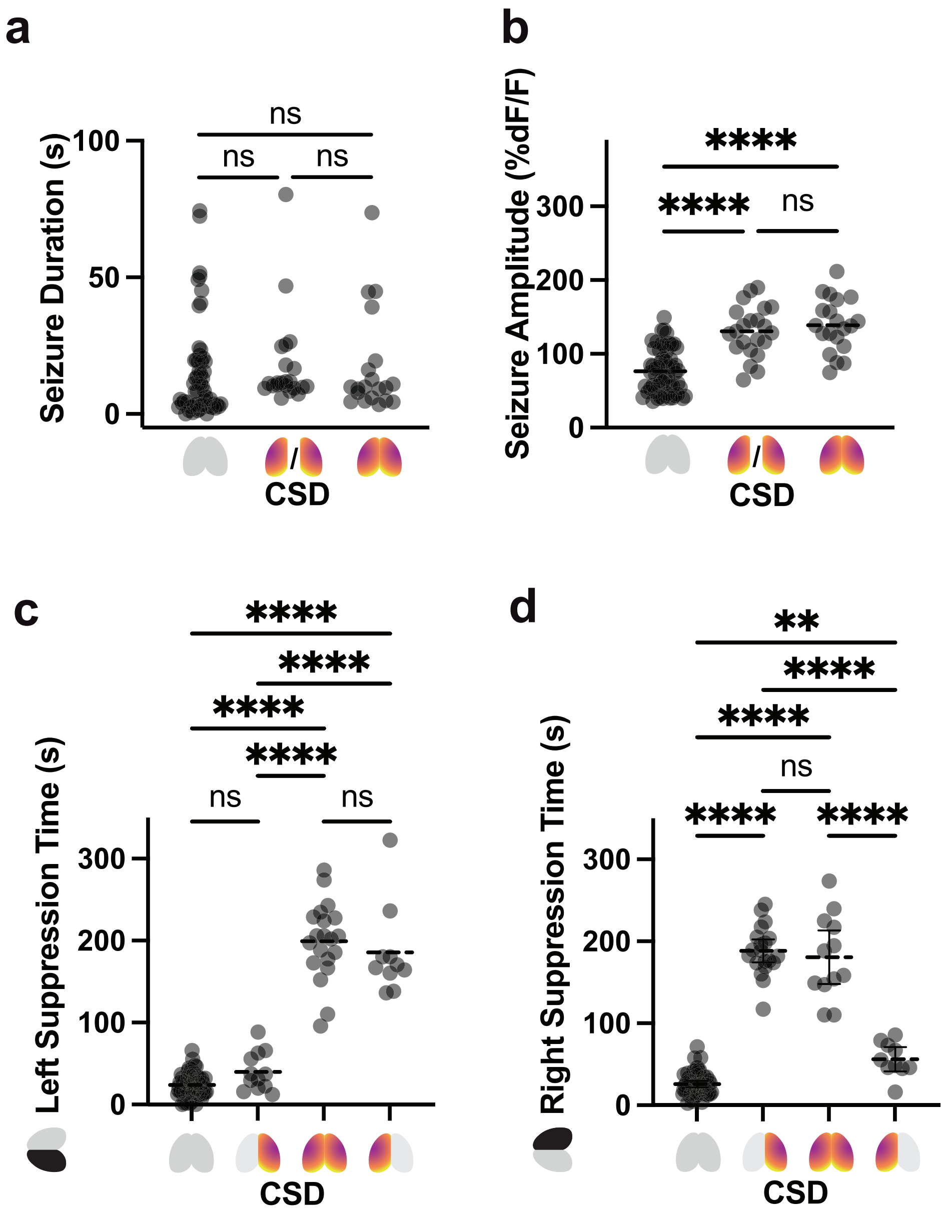


**Extended Data Figure 3 – Summary Metrics of All Seizure and CSD events**

**a,** Average peak fluorescence of seizures that were followed by no CSD, unilateral CSD (pooled right and left), or bilateral CSD. Statistical analysis by Kruskal-Wallis test for non-Gaussian data distribution with Dunn's multiple comparison correction.

**b,** Average duration of seizures that were followed by no CSD, unilateral CSD, or bilateral CSD. Same statistical approach as a. (p**** < 0.0001, α = 0.05).

**c,** Left hemisphere post-event suppression time (time to return of baseline 1-3 Hz slow wave power) after seizure alone (no CSD), right unilateral CSD, left unilateral CSD, or bilateral CSD. Statistical analysis by Brown-Forsythe test for normally distributed data, with Dunn's multiple comparison correction (p **** < 0.0001, α = 0.05).

**d,** Right hemisphere post-event suppression time (time to return of baseline 1-3 Hz slow wave power) after seizure alone (no CSD), right unilateral CSD, left unilateral CSD, or bilateral CSD. Statistical approach same as c. (p** < 0.01, p**** < 0.0001, α = 0.05).

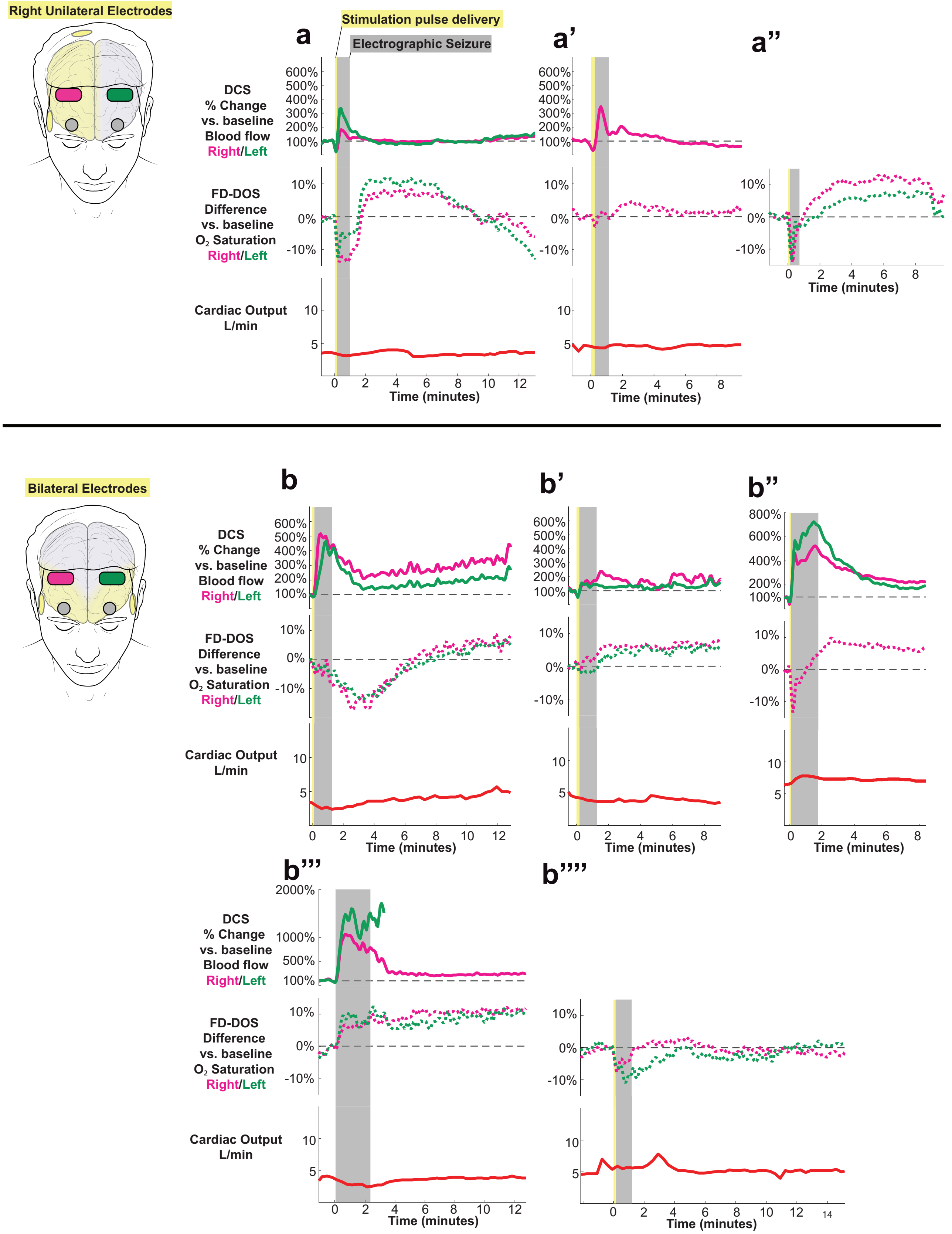

Right Unilateral Electrodes
Stimulation pulse delivery
Electrographic Seizure
a
a'
a''
DCS
% Change
vs. baseline
Blood flow
Right/Left
FD-DOS
Difference
vs. baseline
$O_2$ Saturation
Right/Left
Cardiac Output
L/min
Time (minutes)
Bilateral Electrodes
b
b'
b''
b'''
b''''

**Extended Data Figure 4. Extended SWEET COMBO Data**, including all additional recordings. See Extended Data methods for data quality control and exclusion criteria.
**a,** Three recordings of right unilateral ECT from three separate patients (see Table 2)
**b,** Five recordings of bitemporal ECT from five separate patients. In all recordings except for one (b’’’’), there is clear evidence of post-ictal waves of hyperperfusion evidenced as increased CBF >200% above baseline, or $\Delta O_2$ % saturation greater than 5%.

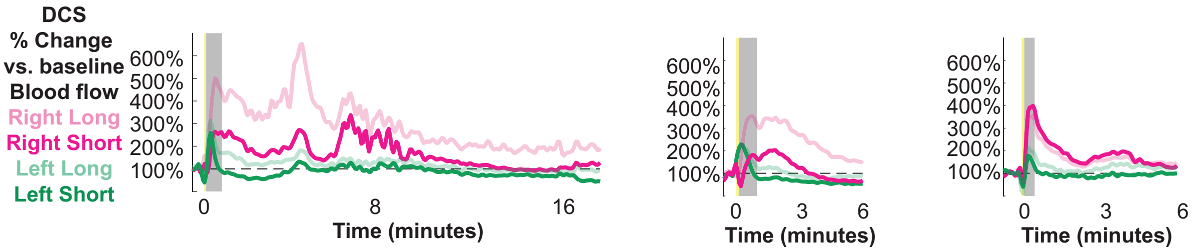


**Extended Data Figure 5.** Right (magenta) and left (green) long source-detector sensor of cerebral blood flow data 4 (transparent lines) overlaid with short source-detector sensor of scalp and skull blood flow (solid lines). Three example recordings presented.

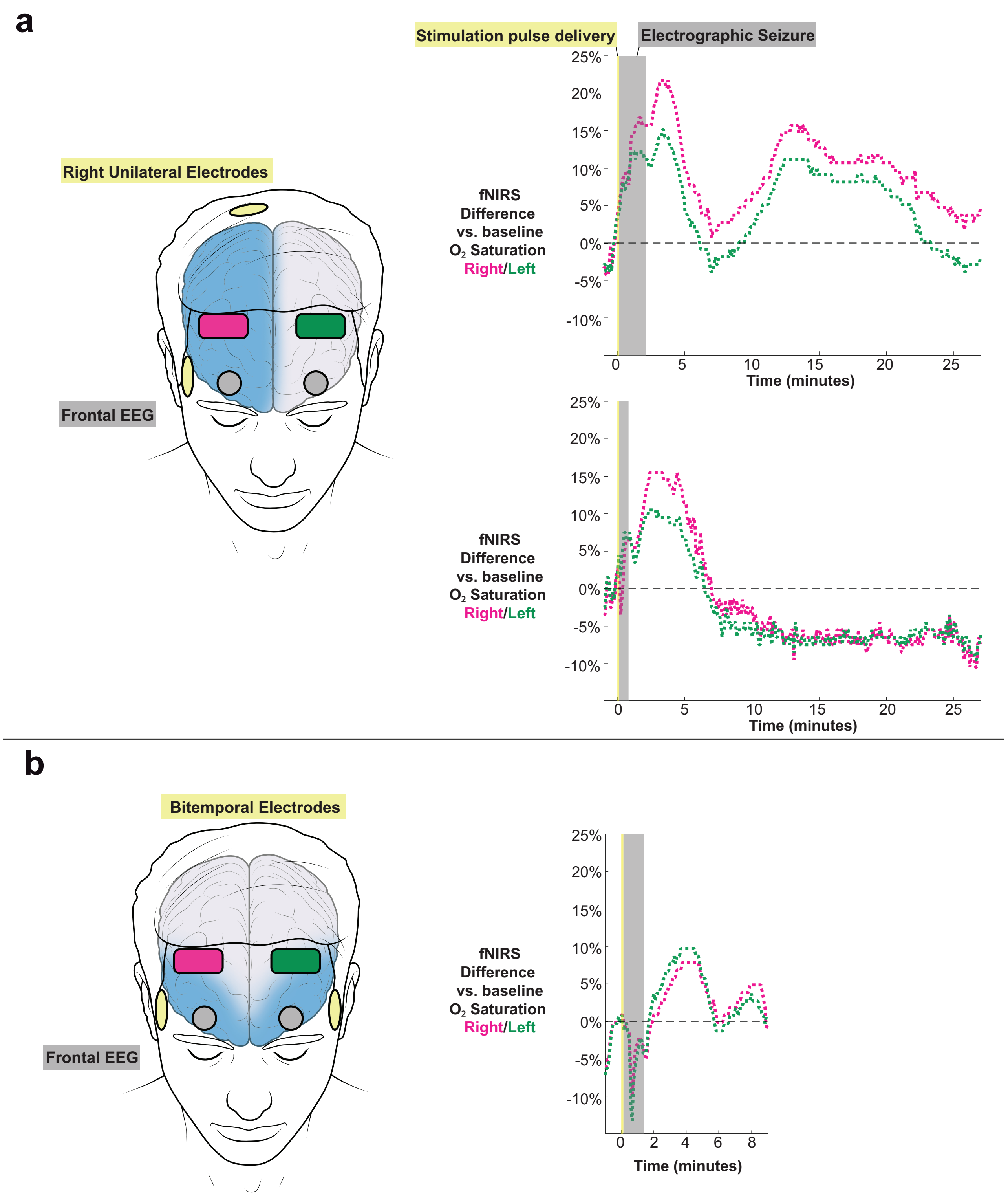


**Extended Data Figure 6.** Case series of human functional near infrared spectroscopy (fNIRS) recordings of brain oxygen saturation during ECT. Two cases of right unilateral ECT, and one case of bitemporal ECT. Data presented as difference from baseline % saturation. Note secondary post-ictal waves of >5% increase from baseline cerebral oxygenation.
