## Supplemental Methods and Discussion for "Electroconvulsive therapy generates a postictal wave of spreading depolarization in mice and humans"

### Supplementary Discussion

All data (**Extended Data Table 2**) were subjected to rigorous quality control, and as a result, n=15 recordings were excluded (see **Supplementary Methods**). Note, in many of the recordings that were excluded for poor signal-to-noise, post-ictal CSD waves were still grossly evident due to their large magnitude. Several variables can potentially lead to poor signal quality and/or introduce confounds in the absolute recovered blood flow index and tissue oxygenation. Motion artifacts can be introduced acutely during electrical stimulation from contraction of the frontalis muscle under the probe, as well as from movement during seizure or repositioning of the head during ventilation. In addition, the electrical stimulus delivery (0.5 – 8 seconds in duration) may induce some degree of thermal heating, though this would be unlikely to persist or fluctuate on the timescale of minutes during subsequent post-ictal CSDs. Other confounds known to impact the absolute hemodynamic metrics include regional variation in tissue optical properties, vascular microanatomy, scalp/skull thickness, head curvature, and hematocrit. *Relative* changes in the cerebral blood flow index and tissue oxygenation, however, are much more robust to these confounds. Accordingly, we standardized analysis across patient recordings by reporting percent change relative to that patient's baseline blood flow index, relative difference from the patient's baseline cerebral oxygen saturation.

Our device uses combined superficial and deep detectors to mitigate against the overlapping effects of skin/skull and brain hemodynamics in the optical path. Somewhat surprisingly, we found that post-ictal surges in blood flow seen by deep cerebral DCS detectors (long source-detector pairs) were mirrored by similar waves in superficial detectors (short source-detector pairs). The relative amplitude of the superficial versus deep post-ictal waves was variable, which we suspect could reflect individual anatomic variations sampled by the tissue light pathlength distribution associated with each probe placement. One possible explanation for this shared signal in brain and scalp/skull is that a massive increase in cerebral blood flow during CSD might exceed vascular autoregulation and spill over collateral flow vessels connecting brain to scalp and skull (e.g., anastomoses from the internal carotid to the supratrochlear or supraorbital arteries); such an effect could potentially cause blood flow changes in superficial detectors. Further studies are required to fully discern the interplay of extra-cerebral and cerebral hemodynamics, which may vary depending on probe placement relative to regional vascular anatomy.

In addition to our experimental results, we described a brief case series (obtained independently at a separate institution) that were consistent with the findings of post-ictal CSD in ECT in both rodent electrophysiology and human brain hemodynamics. In a mouse, post-ictal CSD was observed using a different ECT stimulation paradigm (auricular electrodes), anesthetic agent (isoflurane), and recording modality (glass pipette electrocorticography probe). Seizure induction with ear clip electrodes was notably inconsistent and required relatively higher stimulus intensities. Brief pulse-trains (0.2 s, 60 Hz) induced seizure in 0/3 trials at 10 mA, 0/1 at 30 mA, 4/12 at 50 mA, 2/3 at 70 mA, 11/17 at 80 mA, 8/15 at 90 mA, and 5/7 at 99 mA. In every case, ECT-induced seizures were bilateral, lasting 10-25 seconds. In one case, an ECT-induced bilateral seizure was followed by unilateral SD, confirmed by direct current electrocorticography (**Extended**

**Data Figure 2**). This finding subsequently inspired a human pilot test using a clinical continuous wave NIRS device that reported cerebral oxygen saturation during ECT (**Extended Data Figure 6**), which likewise demonstrated a post-ictal, minutes-long surge in brain oxygen saturation, predominantly in the right hemisphere in two cases of right unilateral ECT, and symmetrically in a case of bilateral ECT.

We further attempted (and ultimately failed) to non-invasively measure the electrical component of CSD waves using state-of-the-art direct current electroencephalography (DC-EEG, ANT Neuro) with both a conventional saline net (n=3) and a custom adaptor with Ag/AgCl electrodes (n=2). We observed slow DC potential drifts during ECT that might be consistent with CSD, but without routine high-pass filters, the low frequency EEG signals were highly sensitive to noise and motion artifacts (e.g., from bag masking ventilatory support). Slow potential drift artifacts could also be generated by small movements in the wires connected to the EEG amplifier (data not shown). Thus, we find that non-invasive DC-EEG is not sufficient to conclusively detect CSD, consistent with prior reports[1,2].

### Supplemental Methods

#### *Diffuse Optical Monitoring Instrumentation and Signal Processing*

The diffuse optical instrumentation employed for this study is a customized, combined frequency domain diffuse optical spectroscopy (FD-DOS) and diffuse correlation spectroscopy (DCS) system (MetaOx, ISS Medical, USA)[3,4]. The FD-DOS system contains diode lasers emitting at four wavelengths (682, 760, 805, and 830nm); the lasers are amplitude modulated at 110MHz. Each of these wavelengths is available at four output ports that operate in a time-multiplexed manner, such that one wavelength is emitting at a time. For detection of the diffusely reflected light, the system contains four photomultiplier tubes. The instrument is constructed in a heterodyne architecture such that the radiofrequency modulated intensity detected by the photomultiplier tubes is down-modulated and compared to a reference signal to recover diffuse light wave amplitude and phase at a rate of 50Hz. The DCS system contains two long coherence length laser sources (DL850-100S, CrystaLaser, USA) operating at 852nm; the laser outputs are attenuated to ~38mW to abide by American National Standards Institute defined limits. For light detection, the DCS system contains two fast single photon avalanche diode (SPAD) arrays comprised of four detectors each (SPCM-AQ4C, Excelitas, CA)[5]. The SPAD array outputs a logic pulse upon detecting a photon that is relayed to a real-time software autocorrelator to recover the normalized intensity autocorrelation function ($g_2(\tau)$) at a rate of 50Hz. The normalized intensity autocorrelation function is used to recover a blood flow index[6].

To perform bilateral cerebral monitoring, two source and two detector ports of the FD-DOS system are dedicated to each hemisphere; for DCS, each hemisphere uses one source and four SPAD detectors. Two identical probes consisting of optical fibers bonded to 90-degree prisms embedded in flexible urethane rubber enable a low-profile, side-firing orientation that can easily adapt to the natural curvature of the skull. Each of the probes contain well-controlled inter-optode source-detector distances that enable optical sampling of tissues at varying depths from the surface. For FD-DOS, these inter-optode distances are 1.5, 2, 2.5, and 3cm, and for DCS, they are 1 and 2.5cm. These distances are selected to balance the increased cerebral sensitivity, expected from larger separations, with sufficient signal-to-noise ratio of the detected light[7]. To enhance the achievable signal-to-noise ratio for DCS, the optical fibers of three SPAD detectors are co-located at the 2.5cm separation to enable signal averaging; only one SPAD is dedicated to the 1cm position. The probes are placed at F3 and F4 on the subject's forehead, positioned laterally to avoid the frontal sinuses, and below the hairline to avoid hair follicle attenuation of the optical signal[8].

Optical properties (absorption and reduced scattering coefficients) and blood flow are extracted via non-linear regression of the measured data using the homogeneous semi-infinite solutions of the frequency domain-photon diffusion equation and the correlation diffusion equation, i.e., for FD-DOS and DCS, respectively[6]. Data from both modalities were down sampled from their original 50Hz collection rate to 0.2Hz to improve measurement signal-to-noise ratio. To further improve accuracy of FD-DOS fitting, data from all inter-optode distances are combined to recover a single set of absorption and reduced scattering coefficients at each wavelength[9]. The absorption coefficients from these fits are then used to recover oxy/deoxy-hemoglobin concentrations using the Beer-Lambert law, while assuming a water fraction of 75%[6]. These concentrations are then used to recover cerebral oxygenation by taking the ratio of oxy-hemoglobin to the sum of oxy and deoxy-hemoglobin concentrations. For DCS, the signals at 1 and 2.5cm are fit independently to preserve the sensitivity to superficial and deep blood flow; the concurrent FD-DOS measurements of tissue absorption and reduced scattering were inputs in the DCS fits. The blood flow indices recovered from the fits are approximately proportional to the average tissue blood flows sampled by the detected light at the 1 and 2.5 cm inter-optode distances. The blood flow index time-series for the 1 and 2.5 cm distances are then divided by their median value in the 30 seconds period prior to stimulation to recover relative blood flow (rBF). Note, probes were calibrated using a silicon phantom with known optical properties before and after each recording to determine FD-DOS light coupling coefficients; specifically, the mean values of phase and amplitude recovered from the silicone calibration block were used to recover the optical coupling coefficients. These coupling coefficients are the multiplicative offset between the measured and theoretically expected amplitude, and the additive offset between the measured and theoretically expected phase.

Determination of whether FD-DOS data provided sufficient quality was made by fitting the multi-distance data to the linearized solutions of the semi-infinite frequency domain-photon diffusion equation[6]. The $R^2$ of the linear fits at each timepoint and each FD-DOS wavelength were recorded for both the amplitude and phase data. If the temporal mean of $R^2$ for the amplitude and/or phase was less than 0.95 for two or more light wavelengths, then the entire subject was excluded from analysis. If this occurred for only one wavelength, the subject was included, but the subject's oxy- and deoxy-hemoglobin concentration were computed using the data from the remaining three wavelengths with mean $R^2 > 0.95$. Finally, for the included subjects and included wavelengths, individual FD-DOS timepoints with $R^2 < 0.95$ for amplitude and/or phase for any of the light wavelengths were removed.

For DCS, datapoints were removed if the photon count rate for a given $g_2(\rho, \tau)$ curve dropped below 10,000 counts per second. Entire subject measurements were removed if photon rates <10,000 counts per second were persistent, or if variation occurred in the quality of $g_2(\tau)$ fits at early decay times, as this would indicate instability in the coherence parameter, $\beta$. This $\beta$ parameter was fit for during a baseline period and subsequently held constant, leaving only blood flow index to be fit for the remainder of the dataset.

*Rodent DC-Electrocorticography Recordings*

All procedures described below were approved by the Institutional Animal Care and Use Committee at the University of New Mexico in compliance with AAALAC guidelines. We measured DC electrophysiologic signals during ECT in n = 7 male C57Bl/6J mice (10 - 12 weeks of age, 27.3 ± 1.0 g). Mice were anesthetized with isoflurane (3% induction, 1.0-1.2 % maintenance, carrier gas $O_2$ 2L/min) and intermittently paralyzed with rocuronium (0.6 mg/kg i.p.) to eliminate muscle artifact. Prior to paralysis, mice were endotracheally intubated and mechanically ventilated (tidal volume 10 - 12 mL/kg, respiratory rate 130 - 140 breaths/min, PEEP 5 cm H20, peak pressure 15 cm H20; Kent Physiosuite, Kent Scientific Corp, Torrington, CT). The scalp was incised and reflected to broadly expose the skull. Bifrontal burr holes were drilled (AP +1.5 mm, ML ±1.5 mm from Bregma), and glass capillary micropipettes (filled with 0.9% NaCl in continuity with Ag/AgCl electrodes) were stereotactically positioned 300 μm beneath the cortical surface. An Ag/AgCl ground electrode was placed in the subcutaneous tissues of the neck. Stimulus trains (12 square-wave pulses, 60 Hz, pulse width 0.2 - 0.5 ms) were generated by a rodent ECT unit (Model 57800, Ugo Basile, Gemonio, Italy) and delivered via auricular clip electrodes moistened with conductive gel. The pulse amplitude was titrated (10 - 99 mA) to identify seizure threshold, then supra-maximal stimuli were delivered up to 99 mA. Unfiltered raw signals were acquired with a DC amplifier (Neuroprobe Model 1600, A-M Systems, Carlsborg, WA) and digitized and recorded at 20 kHz (Power Lab 16/30 and LabChart 7.3.7, AD Instruments, Colorado Springs, CO). Signals were duplicated and digitally filtered off-line to analyze seizure activity (0.5 - 40 Hz band-pass filter) and DC shifts (2 Hz low-pass filter). Total power (0 - 500 Hz) was calculated using a Hann (cosine-bell) FFT of 1024 samples, window overlap 50%.

*Functional Near Infrared Spectroscopy Replication Case Series*

Patients included in this case series (n=3) were recruited from those already receiving ECT at University of New Mexico in 2016. Patients consented to have additional procedural monitoring using the INVOS device (Medtronic, NJ) – a commercially available clinical near-infrared spectroscopy (NIRS) system – to monitor oxygen saturation levels in the right and left hemispheres at the F1 and F2 positions. Note this commercial NIRS system employed for the study is somewhat limited, because it uses a continuous light source. The FD-DOS system in our study modulates the light source intensity to obtain phase and amplitude data that enables separation of absorption and scattering, and thereby leads to more accurate quantification of tissue properties. No additional interventions or changes in patient care were introduced as part of these case series observations. The INVOS instrument's data were sampled at 0.2 Hz to provide continuous monitoring of cerebral oxygenation levels during ECT. Calibration was performed according to the manufacturer's guidelines to ensure accurate and reliable measurements. Data were manually timestamped for stimulation, seizure start, and seizure end time.

*Direct Current Electroencephalography*

A subset of human ECT recordings implemented direct current electroencephalography (DC-EEG) using a portable amplifier (eego, ANT Neuro) combined with either a 24-channel saline net headcap (waveguard, ANT Neuro) with the Cz electrode placed at the skull vertex, or a custom-fabricated adaptor (ANT Neuro) with individual leads coupled to Ag/AgCl electrode cups (WBT-DSC, The Electrode Store) attached to the forehead overlying the bilateral prefrontal cortices using conductive adhesive (Tensive). EEG signals were collected unfiltered and analyzed offline in MatLab using open-source tools (EEGLAB)[11].

*Data Visualization*

Except where specified, color schema used throughout the manuscript were from the viridis family, selected to be perceptually uniform (linear color transitions across scales) and robust to all major forms of colorblindness. We thus used the colormap *viridis* for jRGECO1a fluorescence (%dF/F, yellow-green-blue), *turbo* for pulse current steps (mA, rainbow), and *magma* for pulse frequency groups (Hz, black-purple-red-orange-yellow). Hemoglobin optical intrinsic signal data is zero-centered and thus a diverging *redwhiteblue* colormap was used.

**References:**

1. Hofmeijer, J., van Kaam, C. R., van de Werff, B., Vermeer, S. E., Tjepkema-Cloostermans, M. C. & van Putten, M. Detecting Cortical Spreading Depolarization with Full Band Scalp Electroencephalography: An Illusion? *Front Neurol* **9**, 17, (2018).
2. Riederer, F., Beiersdorf, J., Lang, C., Pirker-Kees, A., Klein, A., Scutelnic, A., Platho-Elwischger, K., Baumgartner, C., Dreier, J. P. & Schankin, C. Signatures of migraine aura in high-density-EEG. *Clin Neurophysiol* **160**, 113-120, (2024).
3. Forti, R. M. *et al.* Non-invasive diffuse optical monitoring of cerebral physiology in an adult swine-model of impact traumatic brain injury. *Biomed Opt Express* **14**, 2432-2448, (2023).
4. Carp, S. A., Farzam, P., Redes, N., Hueber, D. M. & Franceschini, M. A. Combined multi-distance frequency domain and diffuse correlation spectroscopy system with simultaneous data acquisition and real-time analysis. *Biomed Opt Express* **8**, 3993-4006, (2017).
5. *American National Standard for safe use of lasers (ANSI 136.1)*. (The Laser Institute of America).
6. Durduran, T., Choe, R., Baker, W. B. & Yodh, A. G. Diffuse Optics for Tissue Monitoring and Tomography. *Rep Prog Phys* **73**, (2010).
7. Carp, S. A., Robinson, M. B. & Franceschini, M. A. Diffuse correlation spectroscopy: current status and future outlook. *Neurophotonics* **10**, 013509, (2023).
8. Hebden, J. C., Forrester, G., Zhang, H. & Pacis, D. M. Using spectral derivatives to remove the influence of hair on optical images of the static absorbing properties of tissue-like turbid media. *Neurophotonics* **11**, 025002, (2024).
9. Fantini, S., Franceschini, M. A., Fishkin, J. B., Barbieri, B. & Gratton, E. Quantitative determination of the absorption spectra of chromophores in strongly scattering media: a light-emitting-diode based technique. *Appl Opt* **33**, 5204-5213, (1994).

10**.** Boas, D. A. & Yodh, A. G. Spatially varying dynamical properties of turbid media probed with diffusing temporal light correlation. *Journal of the Optical Society of America A* **14**, (1997).
11**.** Delorme, A. & Makeig, S. EEGLAB: an open source toolbox for analysis of single-trial EEG dynamics including independent component analysis. *J Neurosci Methods* **134**, 9-21, (2004).
